# A human tubular aggregate myopathy mutation unmasks STIM1-independent rapid inactivation of Orai1 channels

**DOI:** 10.1101/2022.08.12.503733

**Authors:** Priscilla S.-W. Yeung, Megumi Yamashita, Murali Prakriya

## Abstract

Ca^2+^ release-activated Ca^2+^ (CRAC) channels are activated by direct physical interactions between Orai1, the channel protein, and STIM1, the endoplasmic reticulum Ca^2+^ sensor. A hallmark of CRAC channels is fast Ca^2+^-dependent inactivation (CDI) which provides negative feedback to limit Ca^2+^ entry through CRAC channels. Although STIM1 is thought to be essential for mediating CDI, the molecular mechanism of CDI remains largely unknown. Here, we examined a gain-of-function (GOF) human Orai1 disease mutation, L138F, that causes tubular aggregate myopathy (TAM). Through pairwise mutational analysis, we determine that large amino acid substitutions at either L138 or the neighboring T92 locus evoke highly Ca^2+^-selective currents in the absence of STIM1. We find that the GOF phenotype arises due to steric clash between L138 on TM2 and T92 located on the pore helix. Surprisingly, strongly activating L138 and T92 mutations also show CDI in the absence of STIM1, contradicting prevailing views that STIM1 is required for inactivation. CDI of constitutively open T92W and L138F mutants occurred with similar kinetics as WT Orai1 but showed enhanced intracellular Ca^2+^ sensitivity, which could be normalized by the addition of STIM1. Truncation of the Orai1 C-terminus reduced T92W CDI consistent with a key role for the Orai1 C-terminus for CDI. Overall, these results elucidate the molecular basis of the human TAM-linked mutation and indicate that CDI of CRAC channels is mediated by an Orai1-intrinsic mechanism with STIM1 tuning the calcium sensitivity of CDI.

## Introduction

Store-operated Ca^2+^ release-activated Ca^2+^ (CRAC) channels formed by the Orai1 protein mediate a wide range of Ca^2+^-dependent cellular processes including cell motility, proliferation, and differentiation (Prakriya & Lewis, 2015). Orai1 channels open in response to release of endoplasmic reticulum (ER) Ca^2+^ stores and function as an essential mechanism for Ca^2+^ entry in most cells. The coupling of ER Ca^2+^ stores to the opening of Orai1 channels in the plasma membrane is tightly controlled by the ER Ca^2+^ sensor protein STIM1, which functions as the activating ligand for Orai1 channels (Prakriya & Lewis, 2015). Several types of loss-of-function (LOF) mutations within Orai1 or STIM1 can block channel gating and abrogate store-operated Ca^2+^ entry. In human patients, these defective channels have deadly consequences, causing severe combined immunodeficiency, autoimmunity, and ectodermal dysplasia (Feske, 2010; Lacruz & Feske, 2015), highlighting the essential role of Orai1 channels for immunity and host defense. From a molecular-structural standpoint, Orai channels consist of six subunits forming three layers of transmembrane domains (TMs), with the pore-lining TM1 domains encapsulated by the TM2/3 ring which is in turn surrounded by TM4 helices (Hou *et al*., 2012). Although much of the early attention was focused on the pore-forming TM1 helix, it is increasingly clear that TMs 2-4 are not merely “bystanders” intended for structural support for the pore, but instead play an active and crucial role in relaying the STIM1 binding signal from the peripheral C-terminus to open the channel gate in the central axis (Zhou *et al*., 2016; Frischauf *et al*., 2017; Yeung *et al*., 2018; Yeung *et al*., 2020).

In addition to their high Ca^2+^-selectivity and store-dependent activation, a distinguishing feature of CRAC channels is fast Ca^2+^-dependent inactivation (CDI). CDI was first described in native CRAC currents of T-cells and mast cells (Hoth & Penner, 1993; Zweifach & Lewis, 1995a) and is now known to extend to all Orai isoforms (Orai1-3) when activated by STIM1 (Lis *et al*., 2007). Fast CDI is serves as an important feedback mechanism for restraining Orai channel activity, limiting Ca^2+^ entry at hyperpolarized potentials and preventing cellular Ca^2+^ overload (Zweifach & Lewis, 1995a; Fierro & Parekh, 1999b). Consistent with this feedback role, mutations that limit CDI enhance the frequency of Ca^2+^ oscillations and increase NFAT4 activation in HEK293 cells (Zhang *et al*., 2019). Yet, despite efforts over the last two decades, the molecular mechanism of CDI is not understood. Based on the differential effects of fast and slow Ca^2+^ buffers in T cells and mast cells, CDI is known to arise from Ca^2+^ microdomains around individual Orai1 channels (Zweifach & Lewis, 1995a; Fierro & Parekh, 1999b) with the Ca^2+^ sensing site localized very close to the channel pore estimated to be ∼ 3 nm from the pore (Zweifach & Lewis, 1993). More recent studies have indicated an essential requirement for STIM1 in mediating CDI, with the inhibitory domain (ID) of STIM1 playing a key role in the inactivation process (Mullins *et al*., 2009; Mullins & Lewis, 2016). In particular, STIM1 mutants lacking the ID domain do not show CDI (Mullins *et al*., 2009). In line with the idea that STIM1 plays an essential role in mediating CDI, constitutively active Orai1 mutants such as V102C and P245L do not show CDI in the absence of STIM1, but CDI is restored following co-expression of these mutants with STIM1 (McNally *et al*., 2012; Derler *et al*., 2018). Within the Orai1 itself, structure-function studies have implicated the N-terminus (Bergsmann *et al*., 2011; Mullins *et al*., 2016; Zhang *et al*., 2019), the C-terminus (Lee *et al*., 2009) as well as the intracellular TM2-TM3 loop (Srikanth *et al*., 2010), but the identity of the Ca^2+^ sensor for CDI, the inactivation gate, and the precise role of STIM1 for CDI, including the possibility that it serves as the enigmatic Ca^2+^ sensor, is not resolved.

Although physiologically gated by STIM1, several gain-of-function (GOF) Orai1 mutations have also been recently identified in human patients that cause Orai1 to open independently of STIM1 (Nesin *et al*., 2014; Endo *et al*., 2015; Garibaldi *et al*., 2016; Bohm *et al*., 2017). The ensuing chronically elevated intracellular Ca^2+^ levels in otherwise normally unstimulated cells causes a Stormorken-like syndrome with tubular aggregate myopathy and additional accompanying symptoms specific to the individual mutations (Nesin *et al*., 2014; Endo *et al*., 2015; Garibaldi *et al*., 2016; Bohm *et al*., 2017). Potential pore opening mechanisms of several GOF human mutations have been proposed (Zhang *et al*., 2011; Palty *et al*., 2015; Yamashita *et al*., 2017; Yeung *et al*., 2018; Bulla *et al*., 2019), but currently little is currently known about how the TM2 human mutation, L138F, located at the TM1-TM2/3 ring interface and linked to tubular aggregate myopathy, causes channel activation (Endo *et al*., 2015).

In this study, we addressed the molecular mechanism by which the pathologic Orai1 mutation L138F leads to a constitutively open pore. Our results indicate that L138F causes channel activation due to steric clash with the TM1 residue T92 in the neighboring Orai1 subunit. Consistent with this functional interaction, introduction of large amino acids (Leu, Phe, Trp) at T92 also produce very large, constitutively open Orai1 channels with high Ca^2+^ selectivity. Uniquely, both L138F and T92W channels show CDI in the absence of STIM1, indicating that CDI is an intrinsic feature of Orai1 channels. However, CDI of constitutively active T92W channels exhibits higher intracellular Ca^2+^ sensitivity resulting in high levels of constitutive inactivation at positive holding potentials. This altered Ca^2+^ sensitivity of T92W Orai1 is “normalized” to that of WT Orai1 channels by STIM1. Together, these findings identify a molecular phenotype with broad implications for activation and inactivation of Orai1 channels.

## Results

### The TAM-linked human Orai1 L138F mutation opens the pore via steric clash with TM1

The Orai1 L138F mutation in TM2 was identified using whole exome sequencing in a Japanese family with tubular aggregate myopathy and was found to mediate STIM1-independent Ca^2+^ entry (Endo *et al*., 2015), a finding confirmed by electrophysiological measurements (Frischauf *et al*., 2017). To address the molecular mechanism of L138F Orai1 tonic activation, we overexpressed this mutant in HEK293-H cells in the presence or absence of STIM1 and analyzed Orai1 currents by whole-cell patch-clamping. As previously reported (Endo *et al*., 2015; Frischauf *et al*., 2017), these experiments showed that L138F Orai1 conducts Ca^2+^ in the absence of STIM1. The current was consistent with Orai1 channels based on an inwardly rectifying current-voltage relationship with a reversal potential of 53.5 ± 2.5 mV in 20 mM Ca^2+^ Ringer’s solution, permeation of Na^+^ ions in divalent-free solutions, and blockade by µM concentrations of La^3+^ **(*Figure 1A*)**. Interestingly, unlike many other constitutively open Orai1 mutants (Zhou *et al*., 2016; Frischauf *et al*., 2017; Yeung *et al*., 2018), Orai1 L138F conducts a much larger Na^+^ current compared to its Ca^2+^ current **(*Figure 1A*)** (the ratio of I_Na_/I_Ca_ was 16.1 ± 1.9 (n=13 cells) for L138F compared to 3.5 ± 0.6 (n=10 cells) in WT Orai1 gated by STIM1). Further, following whole-cell break-in, L138F currents exhibited rapid rundown during the first 20-30 seconds **(*Figure 1A*)**. A likely explanation for this feature based on increased steady-state inactivation is discussed later in the paper. Co-expression of STIM1 with L138F resulted in significantly larger currents than those evoked by L138F alone, indicating that STIM1 can substantively enhance and further gate the activity of this mutant following store depletion **(*Figure 1 – figure supplement 1A,B*)**.

**Figure 1.**
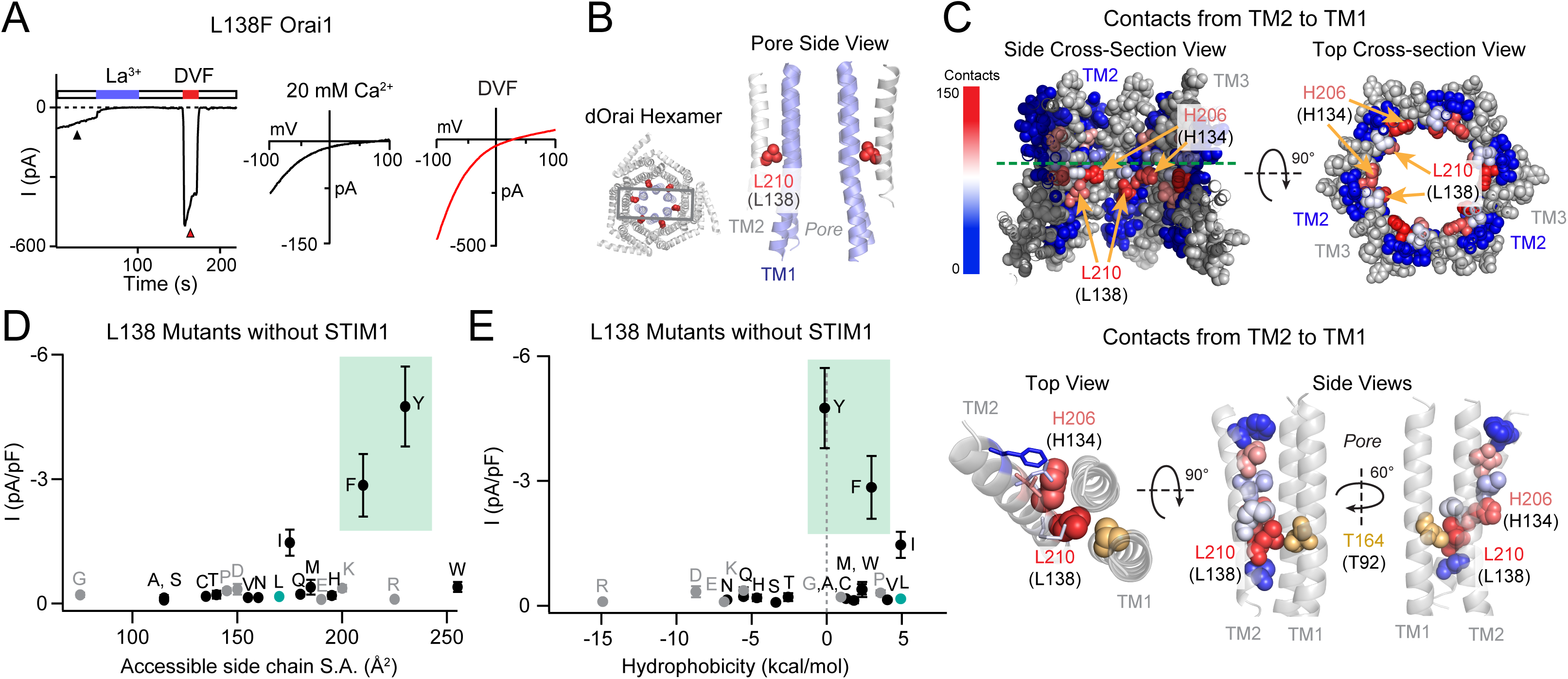
The GOF activity of L138 mutants depends on the size and shape of the introduced side chain. **(A)** Time course of the constitutively active L138F Orai1 current in the absence of STIM1. The trace shows a recording from a HEK293 cell expressing L138F Orai1. At whole-cell break-in (t=0 s), a large standing inward Ca^2+^ current in 20 mM Ca^2+^ external solution is observed which decreases in amplitude over the first 50 s due to current rundown. Replacing the 20 mM Ringer’s solution with a divalent-free (DVF) solution reveals a large Na^+^ current. The current-voltage plots from voltage ramps (−100 mV to +100 mV) are shown on the right. **(B)** Side-view of TM2 residue L210 (hOrai1 L138, red) in the dOrai crystal structure (PDB ID: 4HKR). For simplicity, only two TM1 helices (blue) and two TM2 helices (grey) are shown. Human Orai1 numbering is shown in parentheses. **(C)** Atomic packing analysis of dOrai (PDB ID: 4HKR) showing a heat map of contacts of made by TM2 towards TM1 at the TM2/3 ring-TM1 interface. TM3 is shown in grey and TM4 is hidden for clarity. Residues are colored based on the heat map scale shown on the left. In the bottom panel, TMs 1 and 2 are shown as ribbons, with TM2 residues that face TM1 shown in spheres colored according to the number of contacts made with TM1. TM1 residue T92 is shown in yellow to highlight its proximity to L138. **(D-E)** Current densities of L138 mutants in the absence of STIM1 co-expression plotted against amino acid side-chain surface area and hydrophobicity. L138F/Y channels are open without STIM1 while small and flexible substitutions produce loss-of-function channels. Green shading highlights the GOF mutants L138F and L138Y. N = 4-7 cells. Values are mean ± S.E.M.

How does the L138F mutation cause constitutive channel activity? The crystal structure of *Drosophila melanogaster* Orai (dOrai) in its closed state (PDB ID: 4HKR) shows that L138 located on TM2 protrudes towards the non-pore-lining surface of TM1 (Hou *et al*., 2012) **(*Figure 1B*)**. Atomic packing analysis using a small-probe contact dots protocol to evaluate packing interactions of helical residues (Word *et al*., 1999; Yeung *et al*., 2018) indicated that the majority of L138 contacts are interactions with TM1 **(*Figure 1C*)**. In fact, of the 29 residues located in TM2, L138 displayed the highest number of contacts with TM1 (133 contacts) **(*Figure 1C*)**. H134, a residue critical to channel gating (Frischauf *et al*., 2017; Yeung *et al*., 2018) showed the second highest number of TM1 contacts (125 contacts) **(*Figure 1C*)**. Each of these residues showed substantially more contacts than other residues on TM2, which on average only had 18 contacts with TM1 **(*Figure 1 – source data*)**. This position of L138 and its extensive interactions with TM1 suggested that replacing the native leucine with a bulkier phenylalanine would cause steric clash with TM1 and potentially alter the conformation of TM1.

To test this possibility, we mutated L138 to several other amino acids of different sizes and shapes to assess the contribution of steric effects at this residue. Mutation of L138 to large amino acids with benzene rings such as Phe and Tyr evoked constitutively open channels that conducted ions in the absence of STIM1, whereas mutation of L138 to smaller amino acids did affect L138 activity **(*Figure 1D, see also Figure 1-source data*)**. Hydrophobicity did not seem to be a significant factor in the GOF phenotype **(*Figure 1E*)**. Interestingly, mutation of L138 to His or Trp, which are both larger than Leu, also did not significantly increase baseline Orai1 mutant channel baseline activity **(*Figure 1D*)**, indicating that the channel is more sensitive to a benzene ring than an imidazole ring at this position. These results indicate that the introduction of a large benzene ring at L138 likely leads to steric clash of the exogenous Phe or Tyr side-chains with residues in TM1 causing channel activation. All of the constitutively active L138 mutants showed significantly larger steady-state currents when co-expressed with STIM1, indicating that the constitutively open mutants can be further gated by STIM1 **(*Figure 1 – figure supplement 1*)**.

Intriguingly, analysis of L138 mutant channel activity in the presence of STIM1 co-expression revealed that substitutions of L138 to small amino acids (G/A/S/C/T) caused in loss-of-function mutants that were unable to be activated by STIM1 **(*Figure 1 – figure supplement 1A-C and source data*)**. Mutations of L138 to residues that are polar or charged (N/Q/R/E/K/H), also yielded Orai1 channels with reduced currents in the presence of STIM1 **(*Figure 1 – figure supplement 1A-B and D*)**. The loss of function in these channels is not due to protein misfolding or mistargeting, as measurements of E-FRET between L138A Orai1 and YFP-tagged CRAC activation domain (CAD) showed no loss of CAD binding **(*Figure 1 – figure supplement 1E*)**. Furthermore, L138E Orai1, which could not be gated by STIM1, could be strongly activated by the small molecule modulator, 2-APB, which is known to activate Orai3 and to a smaller extent Orai1 channels in the absence of STIM1 (Yamashita *et al*., 2011), indicating that the L138E mutation disrupts the STIM1-specific gating pathway without affecting channel expression or ability to conduct currents **(*Figure 1 – figure supplement 1D*)**.

Taken together, the phenotypes of the different L138 mutants show that the only substitutions retaining WT store-operated behavior are those that have similar hydrophobicity as the endogenous Leu (V/I/W/M) **(*Figure 1*)**. These results imply that a medium/large hydrophobic residue is needed at position 138 at the TM1-TM2/3 ring interface to correctly relay STIM1-dependent gating signal to the pore. The GOF phenotype of L138F human mutation appears to be driven by steric clash between the introduced benzene ring in Phe with an unknown TM1 residue to evoke channel activation.

### A screen of potential partners on TM1 reveals an L138-T92 interaction

What is the residue on TM1 that clashes with L13F to cause channel activation? In the closed state structure of *Drosophila* Orai (Hou *et al*., 2012), there are three residues on the non-pore-facing side of TM1 within 3 Å of L138 that could act as potential interaction partners for L138. These are: S93 on the same subunit, A94 on the same subunit, and T92 from the neighboring subunit **(*Figure 2A*)** (Hou *et al*., 2012). We hypothesized that L138F likely causes constitutive channel activation via steric hindrance with one of these three residues. To test his hypothesis, we next examined whether relieving this clash by reducing the size of the opposing residue could relieve constitutive channel activity. To test this “rescue” idea, we introduced a Gly at positions 92, 93, and 94 and asked if this reverses the GOF phenotype of L138F Orai1. We found that the A94G substitution did not alter the GOF phenotype of L138F **(*Figure 2B*)**. On the other hand, unlike the L138F single mutant, introduction of Gly at the T92 position in the T92G/L138F double mutant caused the channel to be no longer constitutively active **(*Figure 2B,C*)**. Finally, S93G/L138F channels showed an intermediate phenotype with less activity than L138F but still higher than WT **(*Figure 2B*)**. Together, the results from these rescue experiments suggest that the L138F Orai1 mutant is activated due to steric clash with either S93 on TM1 of the same subunit or T92 of the neighboring subunit.

**Figure 2.**
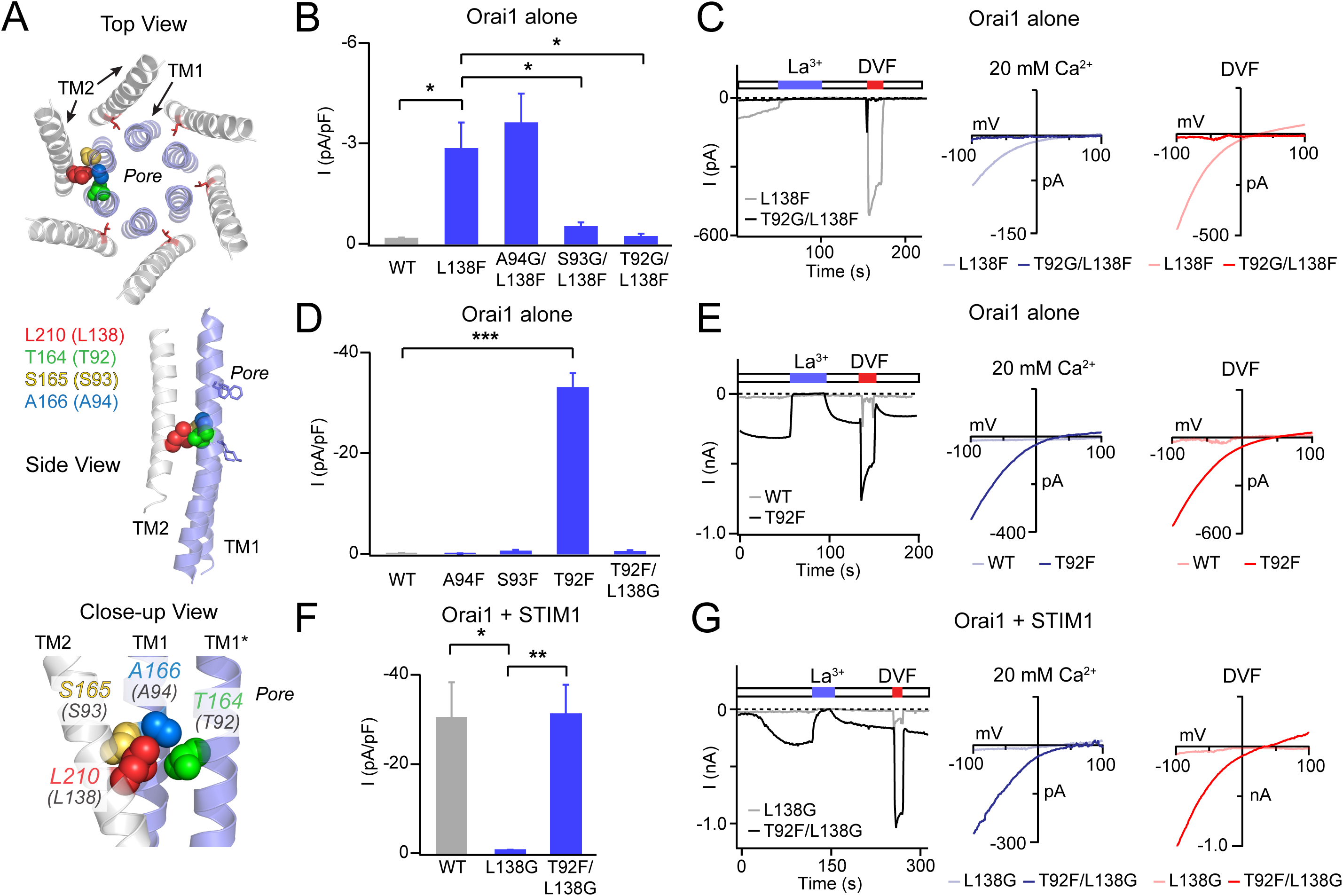
Analysis of L138-TM1 double mutants indicates that L138 interacts with T92. **(A)** Three views of L210 (hOrai1 L138, red) in the dOrai crystal structure (PDB ID: 4HKR) surrounded by the nearby TM1 residue T164 (hOrai1 T92, green) from a neighboring subunit, and S165 (hOrai1 S93, yellow) and A166 (hOrai1 A94, blue) from the same subunit. **(B)** The Orai1 L138F GOF phenotype is reversed by the glycine substitutions at T92 and S93, but not by a Gly substitution at A94. The bar graph plots the current density in the indicated mutants. **(C)** Time course and current-voltage relationships highlighting the reversal of the L138F GOF phenotype by T92G. The L138F single mutant trace and I-V plots are the same as those shown for L138F in Fig. 1A. **(D)** Introduction of a Phe residue at T92, but not at A94 or S93 produces constitutively active GOF channels. This GOF phenotype of T92F is completely reversed by neutralizing the bulky Phe residue at 92 with a Gly substitution at L138. The bar graph shows the current densities in the indicated mutants. **(E)** Example traces of T92F or WT Orai1 currents expressed in the absence of STIM1. T92F evokes a tonically active, Ca^2+^-selective current. **(F-G)** Loss of STIM1-mediated gating of L138G is reversed by a Phe substitution at T92 (L138G/T92F mutant), suggesting that the total size chain size at this locus is crucial for channel activation. N = 4-6 cells. In panels B, D, and F, values are mean ± S.E.M. *p<0.05 by one-way ANOVA followed by unpaired t-test between the indicated variants. **: p<0.01 by unpaired t-test, ***: p<0.001 by unpaired t-test.

If steric clash between L138F and T92 or S93 explains the GOF phenotype of L138F, we reasoned that introduction of large amino acids at these loci in Orai1 should also evoke GOF channels through steric clash with the endogenous L138 residue. We tested this hypothesis by mutating each of these three potential interaction sites on TM1 (A94, S93, and T92) sequentially to Phe, the same amino acid that evokes constitutive activation when introduced at L138 **(*Figure 2D*)**. This analysis indicated that only mutation of T92 to Phe produced constitutively open channels **(*Figure 2D,E*)**. By contrast, S93F and A94F mutant channels were not constitutively active, and in fact these mutations also impeded STIM1-mediated channel function **(*Figure 2 – figure supplement 1*)**. Importantly, as seen with the T92G/L138F rescue experiment, introduction of L138G in the strongly active T92F channel completely reversed its GOF phenotype **(*Figure 2D*)**. Thus, the most straightforward interpretation of these results is that the L138F mutation, and correspondingly, the T92F mutation evoke tonic channel activation due to steric clash between adjacent pairs of TM1 and TM2 helices.

Conversely, because small amino acids at L138 lead to LOF channels that cannot be gated by STIM1, we postulated that this loss of gating could be rescued by restoring the contacts in this region by introducing a larger side-chain at T92. Indeed, the T92F/L138G double mutant shows STIM1-mediated gating **(*Figure 2F,G*)**. Evidently, the steric contacts between the pore helix and TM2 in this region of the channel are finely tuned to relay the STIM1-mediated conformational change to the pore.

### Large amino acids mutations at T92 cause strong activation of Orai1 channels

The results presented above indicate that T92F channels are tonically active due to steric clash between T92F and L138. To address whether the constitutive channel activity of T92F shares side-chain size dependence analogous to that seen at position L138 **(*Figure 1C*)**, we mutated T92 to other large and small amino acids and analyzed the mutant Orai1 channel currents in the absence or presence of STIM1. Consistent with strong side-chain size dependence for the constitutive activity of T92 mutants, we found that substituting larger amino acids at this position produced GOF channels with large Orai1 currents even in the absence of STIM1 co-expression **(*Figure 3A and Figure 3, source data*)**, whereas mutants with small, polar substitutions (G/A/S/C) did not cause constitutive activity but instead retained store-operated behavior **(*Figure 3, Figure 3 – figure supplement 1*)**. In general, amino acids with larger surface area (L/M/F/Y/W) produced much larger Orai1 currents (25-35 pA/pF) compared to amino acids with intermediate size (V/I/H) (5-10 pA/pF) **(*Figure 3A*)**. The largest constitutively active currents were seen with the Orai1 T92L/M/F/Y/W mutations, which produced Ca^2+^-selective currents with reversal potentials approximately 45-55 mV, similar to the highly active Orai1 H134A/C/S/T and ANSGA channels described previously (Yeung *et al*., 2018; Yeung *et al*., 2020). Consistent with the high baseline activity of these mutants, T92L/M/F/Y/W channels were not noticeably further activated following whole-cell break-in when co-expressed with STIM1 **(*Figure 3 – figure supplement 1*)**. Moreover, these T92 mutants displayed much larger currents in 20 mM Ca^2+^ external solution compared to L138F/Y channels, with current densities of 30-35 pA/pF and 3-5 pA/pF respectively **(*Figure 2 and Figure 3*)**. We hypothesize that the substantially larger tonic currents elicited by T92 mutations compared to L138 mutations may be related to the size of the endogenous residue at position 92. Because the side-chain surface area of Thr (140 Å^2^) is smaller than that of Leu (170 Å^2^), replacing T92 with bulky residues likely produces greater steric clash at the TM1-TM2 interface and therefore stronger channel activation than the complimentary L138F substitution, where the native Leu is already quite large. Finally, like the charged substitutions at L138, Asp or Arg substitutions at T92 resulted in loss of function in the presence of STIM1 **(*Figure 3 – figure supplement 1B*)**, indicating that introduction of a charge at this locus inhibits channel gating. Furthermore, as seen in other GOF mutations (Yeung *et al*., 2018), addition of K85E to the N-terminus abrogated T92W currents **(*Figure 3 – figure supplement 1C*)**, indicating an essential requirement of this N-terminus residue for gating all constitutively active mutants.

**Figure 3.**
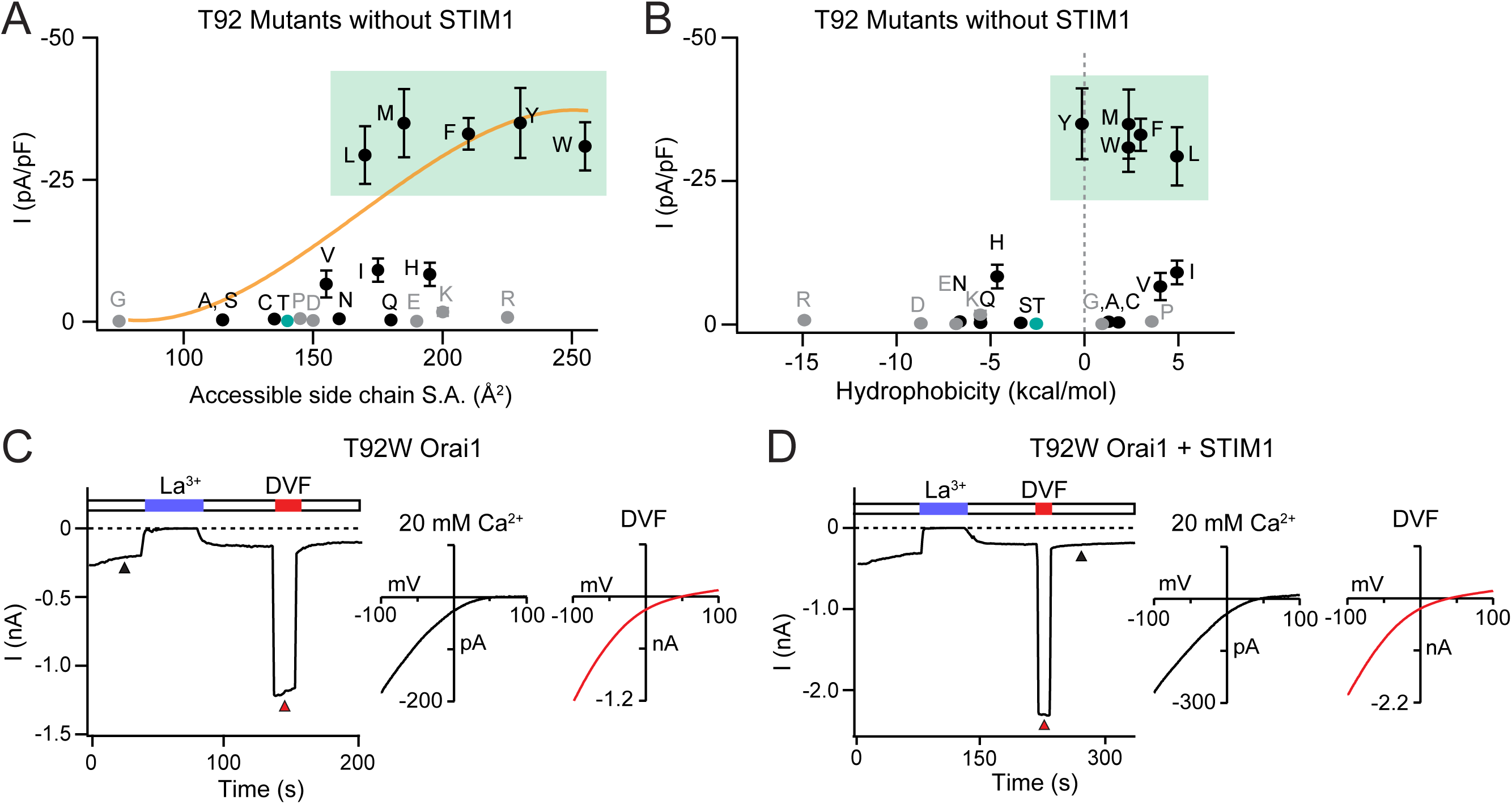
Large amino acid substitutions at T92 cause constitutive Orai1 activation. **(A-B)** The current densities of T92 mutants in the absence of STIM1 co-expression plotted against side chain size and hydrophobicity, showing that bulky substitutions at T92 cause GOF channels. Plots show peak Orai1 currents recorded at −100 mV in HEK293 cells expressing the indicated Orai1 mutants. N = 4-8 cells. Values are mean ± S.E.M. The green shaded areas denote residues that evoke GOF phenotypes. **(C)** Example time course of T92W Orai1 current following whole-cell break-in at t=0 s in a cell expressing T92W Orai1 alone. The I-V graphs in 20 mM Ca^2+^ and DVF solutions are shown the right. T92W Orai1 alone shows inward rectifying I-V curves with positive reversal potentials, similar to STIM1-activated WT Orai1 channels. **(D)** Example time course and I-V graphs of T92W Orai1 with STIM1 co-expression (Orai1:STIM1 = 1:5 cDNA transfection ratio). STIM1 does not significantly boost the current amplitude of GOF T92W Orai1 channels, suggesting that this mutant is nearly fully active at baseline.

### The L138 and T92 GOF mutations exhibit STIM1-independent fast inactivation

Fast Ca^2+^-dependent inactivation (CDI) is a distinguishing feature of CRAC channels in which Ca^2+^ flux through Orai1 at hyperpolarizing potentials (e.g. steps to −100 mV) causes negative feedback inhibition to inactivate the channel on a timescale of tens to hundreds of milliseconds (Hoth & Penner, 1993; Zweifach & Lewis, 1995a; Fierro & Parekh, 1999a; Prakriya & Lewis, 2015) **(*Figure 4A*)**. Fast CDI is regulated by a Ca^2+^-sensing site located very close to the channel pore, estimated to be 3-4 nm away from the pore (Zweifach & Lewis, 1995b) and thought to involve coupling between the inactivation domain (residues 470-491) of STIM1 (Derler *et al*., 2009; Scrimgeour *et al*., 2009; Mullins & Lewis, 2016) with the inner pore (residues 76-91) of Orai1 (Mullins *et al*., 2016). In line with the established requirement for STIM1 for evoking CDI, the GOF Orai1 mutants (including H134A/S, V102C/A, and P245L) do not inactivate without STIM1, but instead show potentiation during hyperpolarizing steps (Zhou *et al*., 2016; Derler *et al*., 2018; Yeung *et al*., 2018) **(*Figure 4 – figure supplement 1B*)**. However, co-expression of STIM1 causes these same mutants to show CDI.

**Figure 4.**
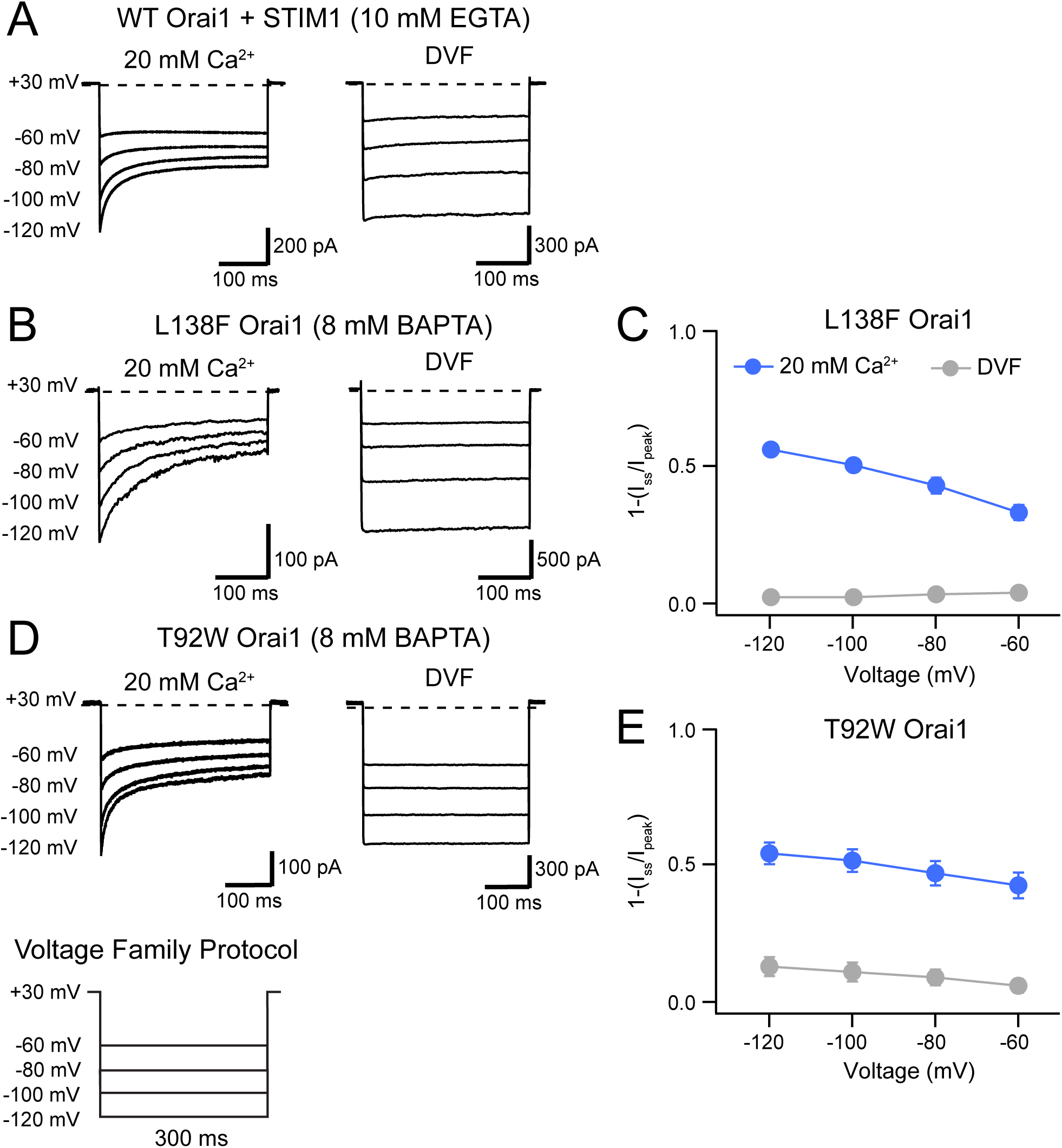
L138F and T92W Orai1 mutants show CDI. **(A)** Currents in response to hyperpolarizing voltage steps in STIM1-gated WT Orai1 channels in the presence of 10 mM EGTA in the internal solution. Hyperpolarizing voltage steps from −120 to −60 mV lasting 300 ms were applied from a holding potential of +30 mV. Replacing the extracellular 20 mM Ca^2+^ Ringer’s solution with a DVF solution eliminates CDI, consistent with the known calcium dependence of CDI (Zweifach & Lewis, 1995a). **(B)** Representative traces showing CDI of L138F Orai1 currents in a cell expressing L138F Orai1 alone. The intracellular solution contained 8 mM BAPTA. **(C)** Quantification of extent of inactivation (1−I_ss_/I_peak_) of L138F GOF mutant channel without STIM1 with 8 mM BAPTA internal solution. **(D)** Representative traces of CDI of T92W Orai1 without STIM1 with an 8 mM BAPTA internal solution. **(E)** Quantification of extent of inactivation (1−I_ss_/I_peak_) of T92W Orai1 in the absence of STIM1. In both mutants, inactivation seen in 20 mM Ca^2+^ is abolished in the presence of divalent-free (DVF) solution indicating that the inactivation requires conducting Ca^2+^ ions. N = 5-17 cells. Values are mean ± S.E.M.

Unexpectedly and in contrast to the behavior of these GOF channels, we observed that several GOF mutants at the L138-T92 gating locus, including L138F/Y and T92F/Y/W, exhibited time-dependent inactivation during hyperpolarizing steps in the absence of STIM1. This is illustrated for the human L138F mutation **(*Figure 4B,C*)**. In the presence of 8 mM intracellular BAPTA, hyperpolarizing steps between −60 to −120 mV caused rapid current decline in the Ca^2+^ currents even in the absence of STIM1 **(*Figure 4B,C*)**, with the current at end of a 300 ms −120 mV voltage pulse declining by ∼60% relative to the peak current. Likewise, expression of T92W alone in the absence of STIM1 co-expression produced Orai1 currents that inactivated during the 300 ms hyperpolarizing step **(*Figure 4E,E*)**. In both mutants, inactivation was strongly reduced or absent in DVF solutions indicating that it is not mediated by Na^+^ ions and suggesting a strong divalent ion-dependence. Consistent with this interpretation, raising extracellular Ca^2+^ concentration from 20 to 110 mM (isotonic Ca^2+^ solution) significantly accelerated and increased the extent of T92W inactivation **(*Figure 4 – figure supplement 1J*)**. Importantly, hyperpolarizing the membrane potential accelerated T92W inactivation and increased its extent, mimicking the effects of raising extracellular Ca^2+^, suggesting that inactivation of T92W strongly depends on the single-channel amplitude.

Two-component fits of the current decay of T92W Orai1 at −100 mV in the 20 mM extracellular Ringer’s solution showed fast and slow τ values of 11 ± 1 ms and 139 ± 19 ms respectively, very similar to kinetics of WT Orai1 currents co-expressed with STIM1 with 10 mM EGTA as the internal buffer (ι−_fast_ = 10 ± 1 ms and ι−_slow_ = 60 ± 8 ms). Thus, the kinetics, divalent ion requirement, and dependence on driving force for Ca^2+^ entry are all features qualitatively similar to CDI of CRAC channels gated by STIM1.

Interestingly, comparison of the extent of inactivation in the different constitutively active L138 and T92 mutants showed that CDI during the hyperpolarizing step is correlated with the size of the introduced side chain **(*Figure 4 – figure supplement 1E-I*)**. Specifically, although T92V was constitutively active, over the course of 100 ms hyperpolarizing steps, this mutant showed no detectable inactivation but instead displayed potentiation similar to other previously described GOF mutations including V102C and H134 (Zhou *et al*., 2016; Derler *et al*., 2018; Yeung *et al*., 2018) **(*Figure 4 – figure supplement 1B*)**. By contrast, larger amino acid substitutions at T92 evoked increasing extent of inactivation with T92W exhibiting the greatest amount of inactivation over the −100 mV hyperpolarizing steps **(*Figure 4 – figure supplement 1I)***. This interesting finding implies that like activation of the GOF L138F and T92W channels, the inactivation may also be induced through steric clash between L138 and T92. To our knowledge, these are the first reported GOF mutants that inactivate in the absence of STIM1.

### Inactivation of T92W Orai1 channels exhibits aberrant dependence on intracellular Ca^2+^ buffering

Previous work has shown that fast CDI is exquisitely sensitive to the species of the intracellular Ca^2+^ buffer (Zweifach & Lewis, 1995a; Fierro & Parekh, 1999b; Yamashita *et al*., 2007). Specifically, fast CDI is not affected by the slow Ca^2+^ chelator, EGTA but is strongly reduced by the fast chelator, BAPTA (Zweifach & Lewis, 1995a; Fierro & Parekh, 1999b), whose *k_on_* of Ca^2+^binding is about 400 times faster than that of EGTA. This is because although both buffers have comparable affinities for Ca^2+^, the slower kinetics of Ca^2+^ binding to EGTA produces a region of relatively unbuffered [Ca^2+^]_i_ around individual CRAC channels. By contrast, the faster *k_on_* of BAPTA ensures that [Ca^2+^] is strongly attenuated in the vicinity (<20 nm) of individual open CRAC channels in this chelator. Therefore, whereas BAPTA is very effective in suppressing both local and global Ca^2+^ signals, EGTA mainly buffers the global Ca^2+^ signal (Neher, 1986). We therefore used the differential effects of EGTA and BAPTA to study the local [Ca^2+^]_i_ dependence of T92W Orai1 inactivation.

In WT Orai1 channels activated by STIM1, we found that reducing intracellular BAPTA from 8 to 0.8 mM increased the extent of CDI **(*Figure 5A*)**, consistent with the known Ca^2+^-dependence of CDI (Zweifach & Lewis, 1995a; Mullins & Lewis, 2016). Likewise, substituting 8 mM BAPTA with the slower chelator EGTA, which should increase local [Ca^2+^]_i_ around CRAC channels, markedly increased the extent of CDI **(*Figure 5A*)**. No significant change in CDI occurred by increasing intracellular EGTA from 10 to 20 mM, consistent with the insensitivity of CDI to changes in the concentration of intracellular EGTA in the mM range (Zweifach & Lewis, 1995a).

**Figure 5.**
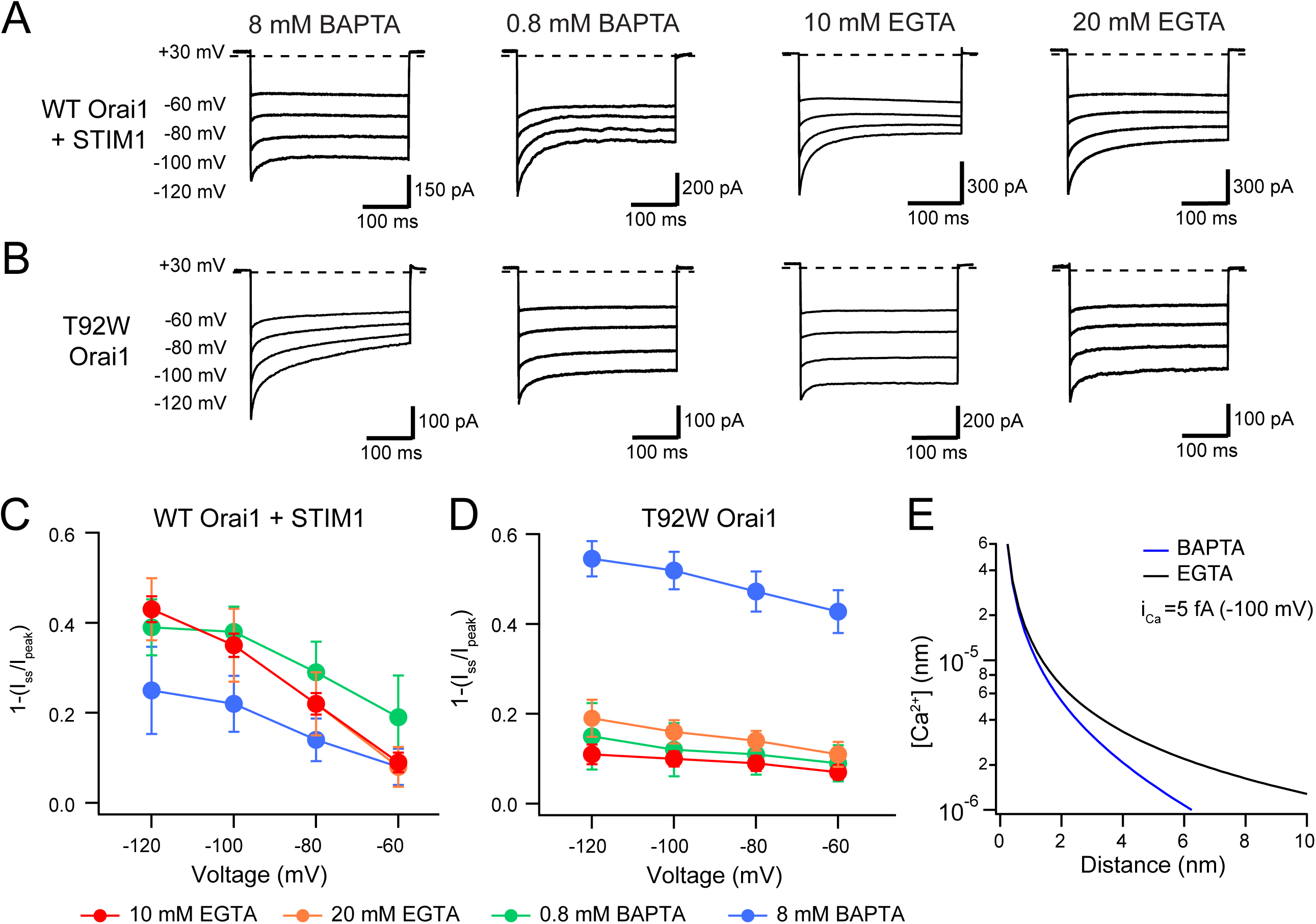
CDI of T92W Orai1 shows enhanced Ca^2+^ sensitivity compared to WT Orai1. **(A-B)** Representative inactivation traces of WT Orai1+STIM1 and constitutively active T92W Orai1 mutant channels during voltage steps with either a slow (EGTA) or fast (BAPTA) Ca^2+^ chelator in the pipette solution. **(A)** In cells expressing WT Orai1, replacing intracellular BAPTA with EGTA or lowering the BAPTA concentration (to 0.8 mM) strongly enhances CDI. **(B)** However, in T92W, replacing BAPTA with EGTA strongly diminishes CDI. Likewise, lowering the BAPTA concentration from 8 to 0.8 mM reduces CDI. **(C-D)** The extent of inactivation (1−I_ss_/I_peak_) plotted against membrane voltage in the indicated buffer solutions. The extracellular Ringer’s solution contained 20 mM Ca^2+^ in all cases. N = 4-17 cells. Values are mean ± S.E.M. Note that the inactivation quantification plot for T92W in 8 mM BAPTA is also shown in Fig. 4E. **(E)** [Ca^2+^]_i_ profiles in the presence of 8 mM BAPTA or 10 mM EGTA. [Ca^2+^] was estimated from equation 1 (see Results). The unitary Ca^2+^ current amplitude was assumed to be 5 fA. At distances beyond 2 nm, [Ca^2+^]_i_ is significantly buffered by BAPTA but not EGTA.

Unexpectedly, fast CDI of T92W showed a very different phenotype in response to changes in the species and concentrations of intracellular buffers. In T92W Orai1, the highest extent of CDI occurred at 8 mM BAPTA and the extent and rate of inactivation at this concentration was broadly comparable to CDI seen in WT Orai1 channels at −120 mV with 10 mM EGTA **(*Figure 5B*)**. Reducing the concentration of intracellular BAPTA to 0.8 mM paradoxically *decreased* CDI in the constitutively active T92W mutant **(*Figure 5B*)**, in contrast to the increase seen in WT Orai1 channels. Most strikingly, replacing the intracellular buffer to 10 mM EGTA caused marked loss of CDI with only a small hint of current decay apparent during the hyperpolarizing steps **(*Figure 5B*)**, in contrast to the behavior of WT Orai1 channels which displayed significant *increases* in CDI in EGTA **(*Figure 5A*)**. Overall, these trends indicated that the pattern of CDI seen with EGTA and BAPTA is essentially reversed in the T92W mutant.

What is the explanation for this aberrant dependence of T92W Orai1 CDI on intracellular Ca^2+^ buffering? We considered and excluded several possibilities. First, we weighed whether the observed inactivation is not Ca^2+^-mediated but instead occurs due to changes in membrane voltage. However, this notion is not consistent with observations indicating that CDI is lost in the presence of Na^+^ as the current carrier, and that raising extracellular Ca^2+^ from 20 to 110 mM Ca^2+^ significantly accelerates and increases the extent of CDI (***Figure 4 – figure supplement 1J***), both of which indicate strong Ca^2+^-dependence for T92W inactivation. A second possibility is that inactivation of T92W Orai1 channels in EGTA containing internal solutions may occur too fast to be detected over the 300 ms hyperpolarizing steps, representing some unknown type of ultra-fast activation. However, careful examination of the initial current immediately following membrane hyperpolarization revealed no such component, and increasing the sampling the current sampling rate from 5 kHz to 20 kHz failed to reveal the presence of an ultra-fast inactivating current that was missed in our standard recording conditions. A third possibility, that T92W disrupts inactivation gating is ruled out by the fact that CDI is very clearly seen in the presence of 8 mM BAPTA and occurs with kinetics similar to that of WT Orai1 channels in EGTA-containing internal solutions. A fourth possibility is that CDI of the T92W still requires STIM1, with the endogenous pool of STIM1 in HEK293 cells interacting with the T92W mutant Orai1 channel with increased affinity to drive CDI. However, T92W current recordings in in STIM1/STIM2 double knock-out HEK293 cells (Emrich *et al*., 2019) also showed continued persistence of CDI, indicating that the CDI of T92W occurs independently of STIM1 and STIM2 **(*Figure 4 – figure supplement 1K*)**. Finally, introducing the E106D Orai1 mutation, which abrogates CDI of heterologously-expressed Orai1 channels (Yamashita *et al*., 2007), abrogated inactivation of T92W Orai1 currents **(*Figure 4 – figure supplement 1L*)**, confirming that the basic mechanism of inactivation gating are shared between T92W and WT Orai1 channels. Together, these results indicate that although the gating mechanism that causes inactivation is likely similar, the Ca^2+^ sensing process upstream of the conformation changes that driven inactivation gating is altered in T92W mutant channels.

### CDI of constitutively active T92W Orai1 exhibits increased Ca^2+^ sensitivity

One clue for why T92W Orai1 shows greater CDI in the presence of BAPTA comes from comparison of the extent of CDI of WT and T92W Orai1 channels in different buffering conditions **(*Figure 6A*)**. The steady-state inactivation plots show that the extent of CDI of WT Orai1 channels at −120 mV in 10 mM EGTA is quantitatively similar to CDI of T92W Orai1 at −60 mV in 8 mM BAPTA **(*Figure 6A*)**. Diffusion models predict that the magnitude of the Ca^2+^ concentration in Ca^2+^ microdomains around individual Orai1 channels should be strongly influenced by the properties of the buffer and the single channel Ca^2+^ current amplitude (*i_Ca_*) (Neher, 1986; Stern, 1992). Specifically, the steady-state [Ca^2+^] as a function of distance *d* from a point source of Ca^2+^ influx can be given by the relation:

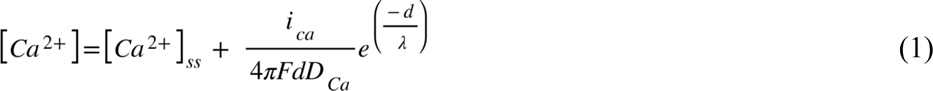

**Figure 6.**
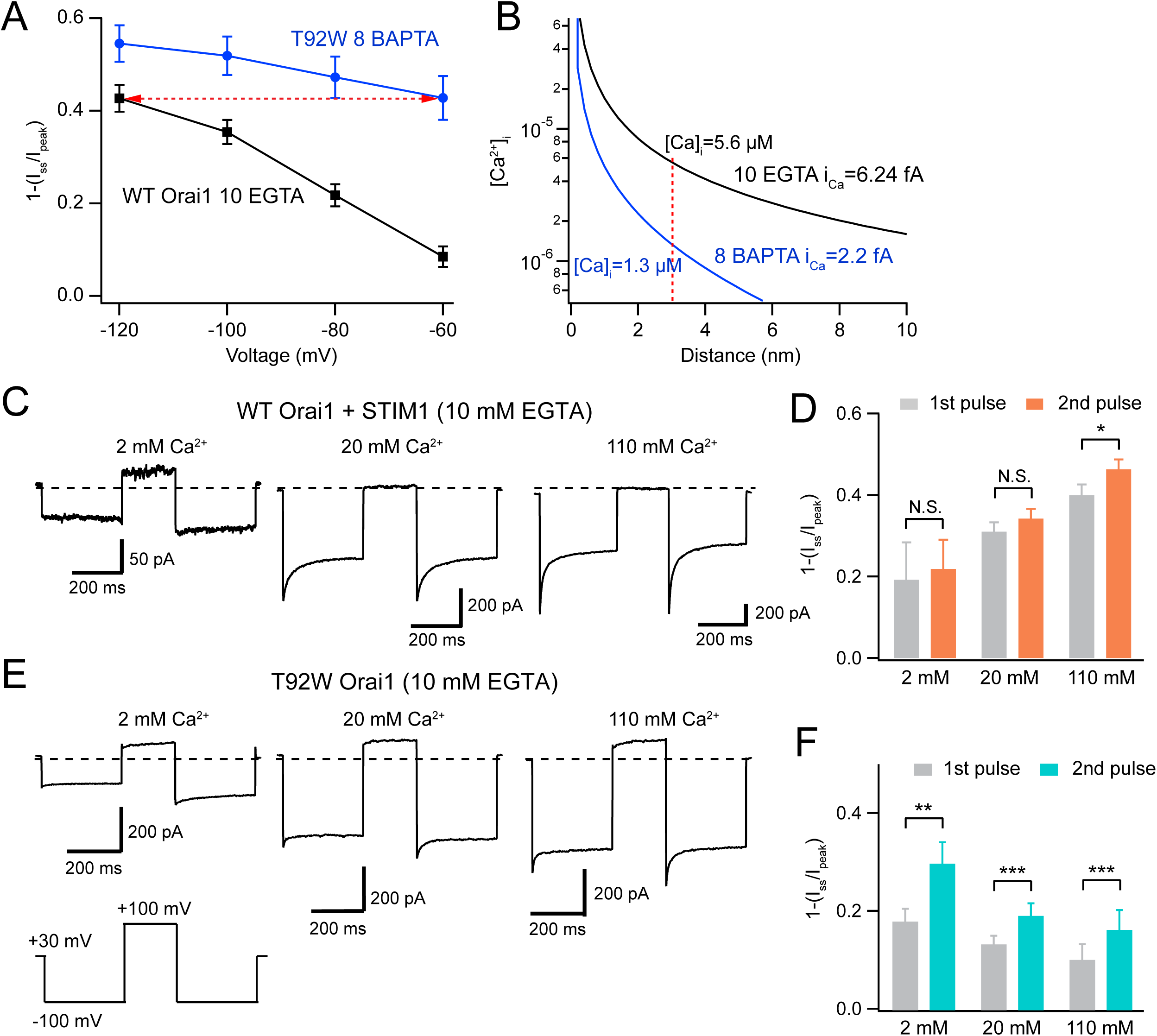
Analysis of recovery from fast inactivation indicates increased Ca^2+^ sensitivity of T92W Orai1 CDI. **(A)** Comparison of extent of CDI in constitutively active T92W Orai1 (in 8 mM BAPTA) and WT Orai1+STIM1 currents (in 10 mM EGTA). The extent of CDI at −60 mV in T92W is identical to CDI in WT Orai1 channels at −120 mV (dashed red arrow). **(B)** [Ca^2+^]_i_ profiles in the two conditions that elicit similar CDI in (A). The unitary Ca^2+^ current at each potential was estimated by scaling the estimated i_Ca_ at −100 mV with the ratio of the peak whole-cell current at the different voltages (see Methods). The profiles indicate that although CDI is similar, the [Ca^2+^]_i_ mediating CDI in T92W Orai1 at −60 mV is substantively lower than the [Ca^2+^]_i_ in WT Orai1 at −120 mV. **(C-F)** Analysis of recovery from inactivation in response to termination of Ca^2+^ influx with depolarizing voltage steps. **(C)** After a −100 mV hyperpolarizing voltage step to induce CDI, the cell was depolarized to +100 mV to abruptly terminate Ca^2+^ influx and lower submembrane [Ca^2+^]. Following the recovery interval at +100 mV (200 ms), a second pulse to − 100 mV was applied to evoke CRAC current. In WT Orai1 channels, the second pulse evokes Orai1 current with similar amplitude and inactivation as the first pulse. Holding potential = +30 mV. **(E)** Recovery from inactivation in T92W Orai1. Following a recovery step to +100 mV to terminate Ca^2+^ influx, a second hyperpolarizing pulse to −100 mV reveals significantly larger peak current and re-appearance of CDI indicating that abruptly terminating Ca^2+^ influx partly restores inactivation of T92W Orai1 channels in 10 mM EGTA. **(D and F)** The extent of inactivation (1 − I_ss_/I_peak_) in the first and second pulses at different extracellular Ca^2+^ concentrations. For T92W channels, CDI was enhanced by pre-pulse with +100 mV in 2 mM, 20 mM, and 110 mM external Ca^2+^ solutions. N = 6-8 cells. Values are mean ± S.E.M. *:p<0.05, **p<0.01, ***: p<0.001 by paired t-test.

where [Ca^2+^]_ss_ is the bulk [Ca^2+^]_i_ (estimated to be negligible in the presence of exogenous buffers), *F* is the Faraday’s constant, *D_Ca_* is the diffusion constant for Ca^2+^ (3×10^−10^ m^2^s^-1^). *λ,* the space constant for Ca^2+^ diffusion in the presence of a buffer (EGTA or BAPTA) at a concentration [B] is given by the relation:

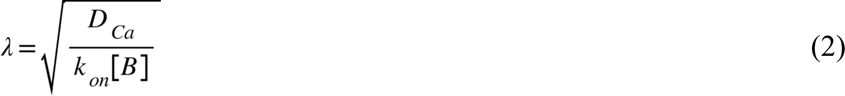

The estimated unitary current of CRAC channels from noise analysis is ∼3.5 fA at –80 mV in 22 mM [Ca^2+^]_o_ (Zweifach & Lewis, 1993). *i_Ca_* is likely about 40% higher due to the high open probability of single CRAC channels during brief (200-300 ms) sweeps (Prakriya & Lewis, 2006), which would be expected to depress current noise. With an estimated *i_Ca_* of 5 fA, the predicted [Ca^2+^]_i_ profiles in BAPTA and EGTA as a function of *d* indicates that [Ca^2+^]_i_ is substantively reduced in 8 mM BAPTA compared to 10 mM EGTA (***Figure 6B***). At less hyperpolarized membrane potentials (−60 to −80 mV), [Ca^2+^] is diminished due to reduction in *i_Ca_*. With the assumption that the number of channels and channel *P_o_* is unchanged by hyperpolarizing steps, we calculated *i_Ca_* at each potential by scaling the unitary current estimate by the ratio of the peak current at each potential to the current at −100 mV. At a distance of 3 nm (estimated to be the distance of the inactivation binding site from the CRAC channel pore (Zweifach & Lewis, 1995a)), these estimates of *i_Ca_* predict that [Ca^2+^]_i_ is ∼1.3 µM at a membrane voltage of −60 mV and 8 mM BAPTA. By contrast, in 10 mM EGTA and at −120 mV, local [Ca^2+^]_i_ is ∼5.6 µM (***Figure 6B***). Thus, the similarity of inactivation of T92W in 8 mM BAPTA at −60 mV to that of WT Orai1 in 10 mM EGTA at −120 mV (***Figure 6A***) indicates that the Ca^2+^-sensitivity of inactivation of T92W Orai1 is substantively increased, such that constitutively active T92W currents inactivate at lower concentrations of intracellular Ca^2+^ compared to WT Orai1 channels (***Figure 6B***). Note that this conclusion is not dependent on the exact values of *i_Ca_* or the precise distance of the Ca^2+^ binding site from the pore, for while altering these parameters would be expected to change the absolute values of [Ca^2+^] at the putative Ca^2+^ binding site, the greater inactivation of T92W channels in BAPTA containing solutions still indicates that these channels show CDI at lower levels of [Ca^2+^]_i_ than WT Orai1 channels.

What are the implications of the increased Ca^2+^-sensitivity of inactivation in T92W Orai1 channels for macroscopic Orai1 currents? We analyzed this question in terms of the simple reaction scheme outlined below.

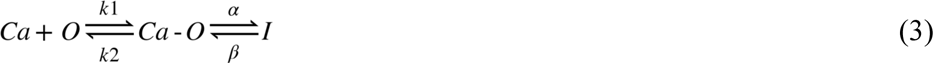

where Ca^2+^ ions bind to open (*O*) CRAC channels with forward and reverse rate constants of *k_1_* and *k_2_* and drive them into the inactivated state (*I*) with forward and reverse rate constants of *⍺* and *β* respectively. Purely for simplicity, we have assumed first-order kinetics for the binding and gating steps, although the actual state diagram is likely more complex (as already hinted by the presence of at least two exponentials for the inactivation time course). Nevertheless, the simple scheme above is instructive in illuminating the potential mechanisms of the change in CDI in T92W Orai1 channels. The analysis of Figure 6 suggests that the Ca^2+^ sensitivity of T92W Orai1 for CDI is strongly enhanced compared to WT Orai1 channels (i.e. *K_d_* =*k_2_*/*k_1_* is reduced in T92W Orai1). This change in *K_d_* predicts that at a given [Ca^2+^]_i_, the forward reaction will be more strongly favored in the mutant compared to WT Orai1, and the overall reaction is predicted to reach equilibrium at lower Ca^2+^ concentrations in T92W compared to WT Orai1 channels. In fact, with weak Ca^2+^ buffering (EGTA or low concentrations of BAPTA) and recurring (every 1 s) steps to −100 mV, T92W inactivation could reach equilibrium at the holding potential (+30 mV) itself such that hyperpolarizing steps are unable to cause additional inactivation. Thus, the most straightforward interpretation of the results is that T92W Orai1 increases the Ca^2+^ sensitivity of CDI of the constitutively active channel. As a result, mutant channels reach equilibrium for CDI at the holding potential (+30 mV) itself in EGTA-containing internal solutions and membrane hyperpolarization fails to elicit additional inactivation.

### CDI of T92W Orai1 in EGTA is unmasked by rapid termination of Ca^2+^ influx

If CDI of T92W Orai1 is at equilibrium due to the steady-state submembrane [Ca^2+^]_i_ at the holding potential (+30 mV), then abruptly lowering submembrane [Ca^2+^]_i_ in EGTA containing solutions should at least partly restore inactivation of T92W Orai1 by moving the reaction leftwards, thereby supporting re-entry of channels from *O* to *I* states during subsequent hyperpolarizing steps. We examined this idea using a two-pulse protocol in which we delivered a +100 mV depolarizing step between two hyperpolarizing pulses to −100 mV. We reasoned that a depolarizing step to +100 mV should abruptly terminate most Ca^2+^ influx. Subsequent chelation of submembrane Ca^2+^ by EGTA should promote recovery of T92W Orai1 channels from CDI. Zweifach and Lewis (Zweifach & Lewis, 1995a) have previously shown that recovery from CDI proceeds with a time course of tens of milliseconds, with ∼90% recovery occurring in 200 ms at a holding potential of −12 mV. Therefore, we reasoned that a 200 ms depolarizing step to +100 mV between two hyperpolarizing steps should promote recovery of Orai1 from CDI.

We tested this idea using a paired-pulse protocol (to −100 mV to evoke I_CRAC_) interspersed with a 200 ms step to +100 mV to abrogate Ca^2+^ influx **(*Figure 6C-F*)**. We hypothesize that if submembrane Ca^2+^ in the +30 mV condition is causing steady-state inactivation in a population of channels to reduce the amount of inactivation that is elicited during the hyperpolarization step, then the 200 ms to +100 mV step before the second test step should reduce the submembrane Ca^2+^ present and therefore enhance the apparent inactivation measured. Using this dual hyperpolarization voltage-clamp protocol, we assessed recovery of T92W Orai1 current inactivation at three extracellular Ca^2+^ concentrations (2 mM, 20 mM, and 110 mM) applied to the same cells in the presence of 10 mM intracellular EGTA. Depolarizing the membrane potential to +100 mV in 2 mM extracellular Ca^2+^ resulted in strong enhancement of the inward current in the second pulse **(*Figure 6E,F*)**. Elevating [Ca^2+^]_o_ to 20 mM elicited smaller current enhancement of the second pulse relative to the first pulse, and raising [Ca^2+^]_o_ to 110 mM elicited even less recovery **(*Figure 6E,F*)**. This result suggests that depolarizing pulses to +100 mV lower [Ca^2+^]_i_ and drives reaction *3* leftwards, thereby allowing channels to recover from inactivation, and therefore, showing greater CDI during the second −100 mV pulse. The extent of current recovery, is however, dependent on the extracellular Ca^2+^ concentration with higher Ca^2+^ concentrations causing less current recovery, as would be expected for a process involving Ca^2+^-dependent accumulation of inactivation. These results indicate that T92 Orai1 channels are significantly inactivated in the presence of 10 mM EGTA and that recovery from inactivation is promoted by lowering the submembrane [Ca^2+^]_i_. Taken together, these observations indicate that the intracellular Ca^2+^ dependence of T92W Orai1 channels is strongly sensitized relative to WT Orai1 channels, causing the channels to attain steady-state inactivation at the holding potential (+30 mV) and blocking further inactivation during hyperpolarizing steps under weak buffering conditions.

### C- and N-terminal mutations differentially affect T92W Orai1 inactivation

We next turned our attention to address the molecular determinants of CDI of T92W channels. Previous studies have implicated several domains of the CRAC channel for CDI, including the ID region of STIM1, the Orai1 N-terminus and the Orai2/3 C-terminus (Lee *et al*., 2009; Mullins *et al*., 2009; Mullins & Lewis, 2016). To determine the role of the Orai1 N- and C-termini for T92W CDI, we tested the effects of mutations in these domains on T92W Orai1 CDI. A previous study employing domain swap and site-directed mutations in Orai2 and Orai3 implicated the C-terminus as a key locus regulating Orai2/3 CDI (Lee *et al*., 2009). To address a potential role of the Orai1 C-terminus for T92W Orai1 inactivation, we therefore truncated the Orai1 C-terminus (***Figure 7A***). We found that deletion of the Orai1 C-terminus (Δ267-301) had no effect the constitutive activity of Orai1 T92W currents (***Figure 7B***), indicating that the C-terminus is not necessary for the constitutive gating of this open mutant and reaffirming the STIM-independence of the activity of this mutant. However, T92W Δ267-301 Orai1 currents showed markedly reduced inactivation **(*Figure 7B,C*)** with the extent of inactivation decreasing from ∼50% to ∼25% at −100 mV. This finding indicates that the Orai1 C-terminus is a key determinant of CDI of T92W Orai1 mutant. The presence of residual inactivation (20-30%) in the T92W Δ267-301 mutant however, implies that additional regions outside of the Orai1 C-terminus also make contributions to CDI.

**Figure 7.**
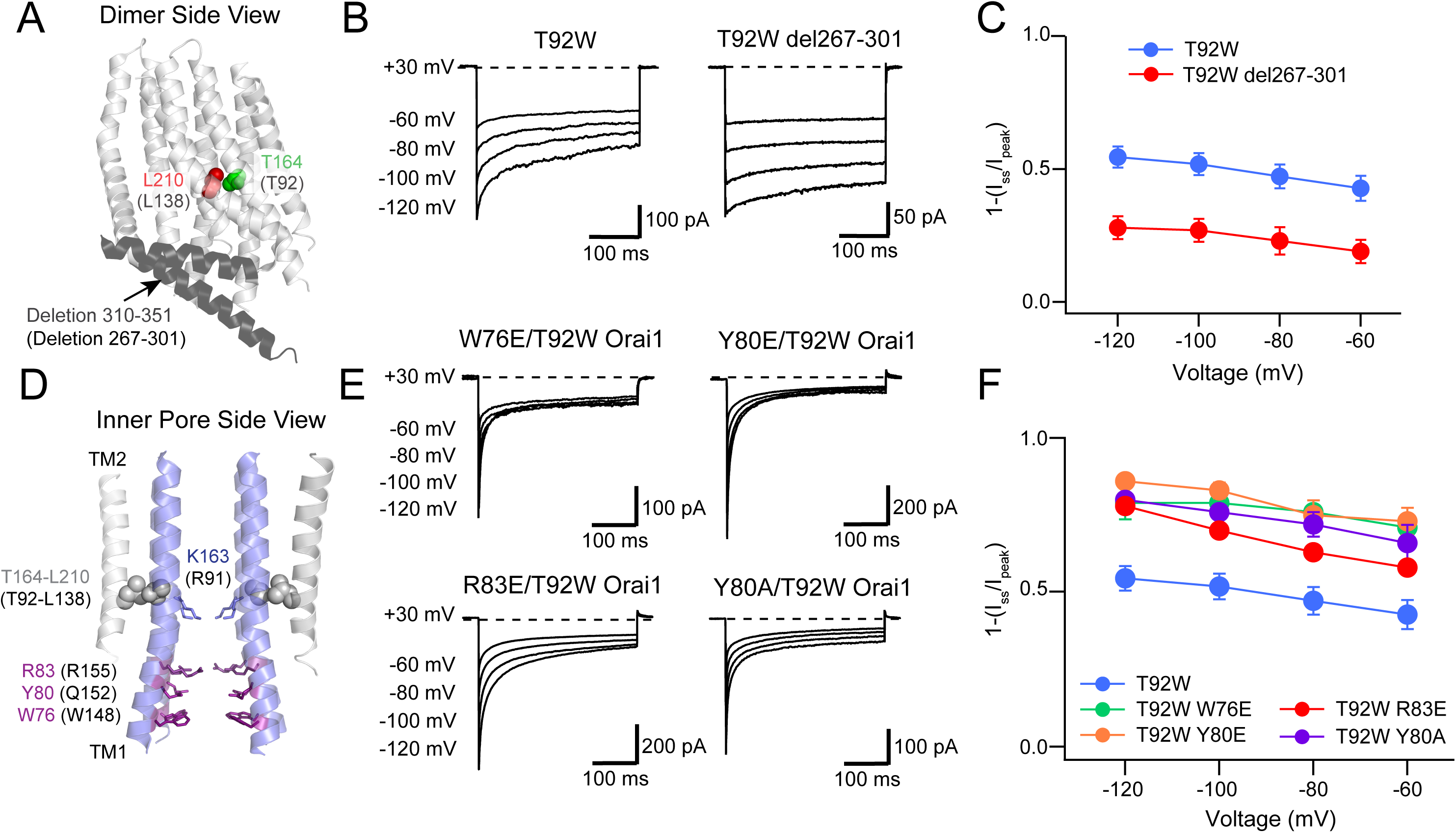
The Orai1 C-terminus contributes to T92W Orai1 CDI. **(A)** Side view of dOrai dimer (PDB ID: 4HKR; hOrai1 numbering in parentheses) showing location of T92 and L138 residues relative to the deleted C-terminal residues 267-301. The Orai1 C-terminus is shown in dark grey. **(B-C)** Deletion of the Orai1 C-terminus residues attenuates inactivation of T92W Orai1. Example traces showing CDI of T92W Orai1 and T92WΔ267-301 Orai1 mutant (B). The extent of inactivation is summarized in panel C. **(D)** Location of N-terminal residues (W76, Y80 and R83) previously implicated in regulating CDI in Orai1 channels. **(E)** Mutations of W76, Y80 and R83 in T92W Orai1 all strongly accelerated CDI compared to T92W Orai1 alone. **(F)** The extent of CDI plotted against membrane voltage in the indicated mutants. The inactivation quantification plot for T92W in (C) and (F) is the same as the data for this mutant in Fig. 4E. N = 4-17 cells. Values are mean ± S.E.M.

At the Orai1 N-terminus, several reports have described changes in CDI in N-terminal Orai1 mutants (Bergsmann *et al*., 2011; Mullins *et al*., 2016; Zhang *et al*., 2019). Specifically, Mullins et al. have shown that mutation of the aromatic residues Y80 and W76 at the most intracellular end of the N-terminal helix can enhance or abrogate CDI in Orai1 channels (Mullins & Lewis, 2016; Mullins *et al*., 2016). Mutagenesis analysis has also implicated three positively charged residues (R91, K87, and R83) located in the inner pore in the TM1 extension helix (Mullins *et al*., 2016). It has been postulated that these residues together with the aromatics referenced above may comprise the inactivation gate or at least involved in the conformational changes mediating CDI (Mullins & Lewis, 2016; Mullins *et al*., 2016). To examine the role of these residues for T92W inactivation, we mutated Y80, R83, and W76 and studied the consequences for T92W CDI. Mullins et al. showed that the Y80A mutation markedly enhances the rate and extent of CDI of Orai1 channels activated by STIM1 (Mullins *et al*., 2016). In a similar fashion, we found that constitutively active Y80A/T92W Orai1 currents showed markedly faster and greater inactivation compared to T92W Orai1 channels (***Figure 7E,F***). Surprisingly, however, the Y80E mutation, which previously was found to abrogate Orai1 CDI, also strongly enhanced the rate and extent of T92W inactivation (***Figure 7E,F***). Similarly, both the W76E and the R83E mutations, which were previously shown to eliminate CDI of WT Orai1 (Mullins *et al*., 2016), accelerated the rate and increased the extent of CDI of T92W Orai1 currents (***Figure 7E,F***). These results are consistent with previous models indicating that the Orai1 N-terminus has an important role in inactivation gating. However, the enhancement of T92W Orai1 inactivation gating by mutations (Y80E, R83E, W76E) that were previously shown to abrogate CDI of WT Orai1 channels reveals a degree of complexity in the role of these residue in controlling inactivation and argue that the aromatics Y80 and W76 regulate inactivation of Orai1 channels through a mechanism distinct from serving as the inactivation gate.

### STIM1 normalizes the inactivation of T92W Orai1 channels

STIM1 is required for CDI of CRAC channels and is postulated to promote inactivation via functional coupling between the ID region of STIM1 with the Orai1 N-terminus (Mullins & Lewis, 2016). What effect, if any, does STIM1 have on the intrinsic CDI of constitutively active T92W Orai1 channels? We examined this question by co-expressing STIM1 together with T92W Orai1 at a STIM/Orai1 cDNA ratio of 5:1, which is expected to be sufficient for STIM1-mediated inactivation of WT Orai1 channels. To our surprise, in the presence of STIM1, the EGTA/BAPTA buffer-dependence of T92W Orai1 CDI was reversed. Currents of T92W Orai1 with STIM1 showed robust inactivation in the presence of 10 mM EGTA comparable in extent and rate to WT Orai1 channels gated by STIM1 **(*Figure 8C,D*)**. Conversely, replacing the intracellular buffer with 8 mM BAPTA (high buffering) strongly reduced CDI of T92W Orai1 channels **(*Figure 8C,E*)**. As summarized in the steady-state inactivation plots, the behavior of T92W Orai1 channels co-expressing STIM1 was essentially comparable to that of WT Orai1 channels activated by STIM1 **(*Figure 8D,E*)**, and fully reversed from the behavior of STIM1-free T92W Orai1 channels. These results indicate that STIM1 “normalizes” the aberrant buffer dependence of inactivation of the constitutively active T92W Orai1 channels. Likewise, co-expressing STIM1 with constitutively active Y80E/T92W Orai1 mutant, which exhibits faster inactivation than T92W Orai1 single mutant in BAPTA containing solutions, also markedly decreased the rate and extent of CDI of this mutant relative to cells without STIM1 co-expression (***Figure 8 – figure supplement 1***). These findings indicate that STIM1 modulates the buffer dependence of CDI of T92W Orai1 channels to make their Ca^2+^ sensitivity similar to that of WT Orai1 channels gated by STIM1. The normalization of CDI by STIM1 is reminiscent of the “normalization” of Ca^2+^ selectivity of V102C and other less Ca^2+^-selective constitutively active Orai1 mutants by STIM1 (McNally *et al*., 2012) and reaffirms the viewpoint that STIM1 plays an essential role for multiple aspects of Orai1 gating including activation, permeation, and inactivation.

**Figure 8.**
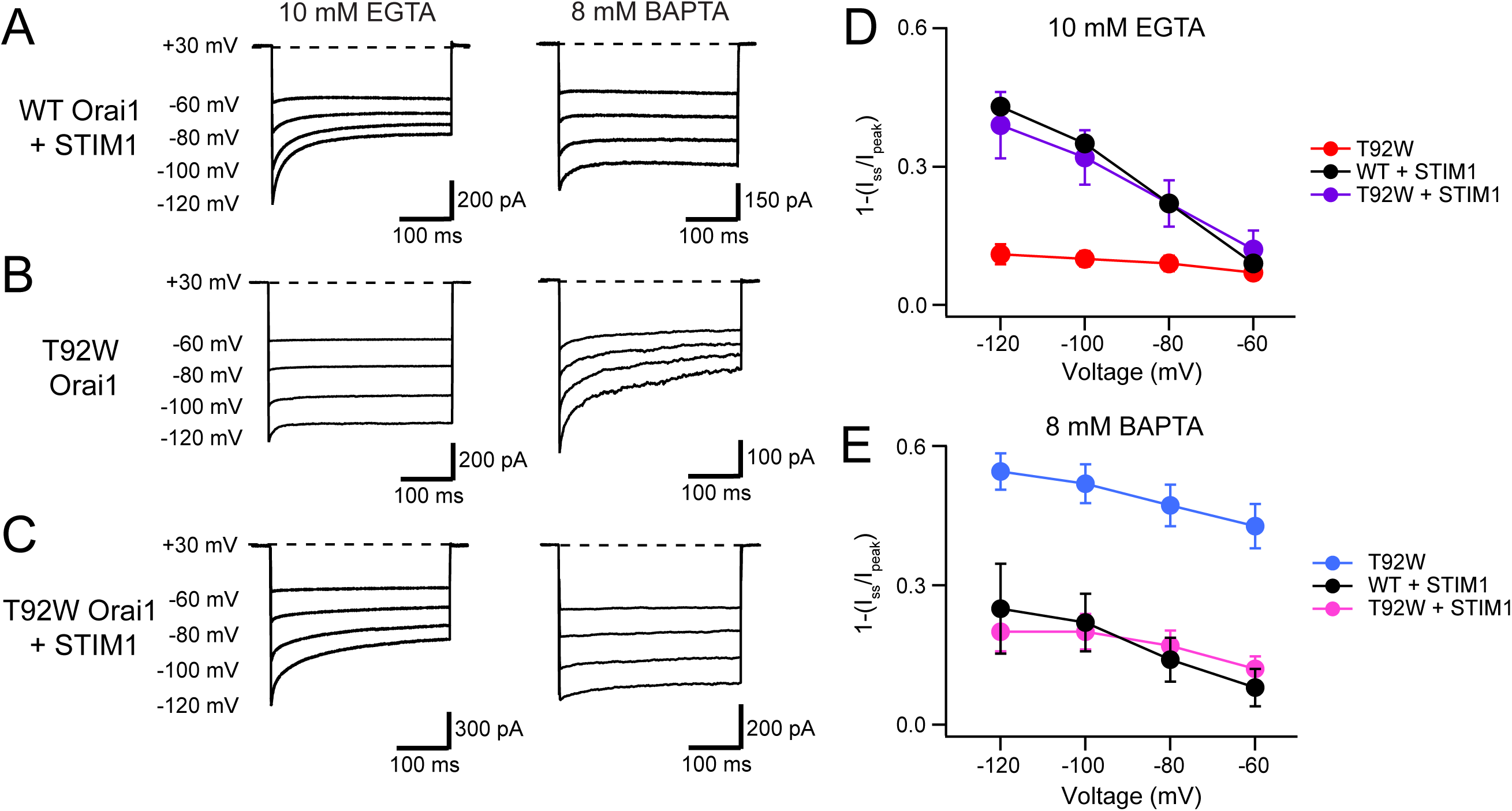
STIM1 modulates the Ca^2+^ sensitivity of T92W Orai1 CDI. **(A-B)** Representative traces of CDI in WT Orai1+STIM1 (A) and T92W Orai1 (B) currents in the presence of EGTA (10 mM) or BAPTA (8 mM). **(C)** The addition of STIM1 normalizes the aberrant intracellular buffer dependence of T92W Orai1 CDI. In the presence of STIM1, CDI of T92 Orai1 is strongly enhanced in 10 mM EGTA compared to T92W currents in the absence of STIM1. Conversely, CDI in STIM1-expressing T92W Orai1 cells in 8 mM BAPTA is reduced relative to cells expressing T92W Orai1 alone (compare right traces in T92W in panels B and C), analogous to the behavior seen in WT Orai1 (A). **(D-E)** Summary plots showing the extent of CDI in the indicated conditions. The inactivation quantification plots for WT and T92W Orai1 are also shown in Fig. 5 C-D. N = 4-17 cells. Values are mean ± S.E.M.

## Discussion

A recent report showed that human mutation Orai1 L138F causes tubular aggregate myopathy with hypocalcemia in human patients (Endo *et al*., 2015). This syndrome is driven by dysregulated Ca^2+^ signaling in muscle cells secondary to constitutive Ca^2+^ entry through open Orai1 channels (Endo *et al*., 2015). In this study, we sought to understand the underlying mechanism of this mutation and determined that steric clash between L138F and T92 on TM1 drives constitutive Orai1 activation. Large amino acid substitutions at either L138 or T92 that increase the amount of contact between these two residues cause GOF Orai1 channels by disrupting the closed state of the pore. By contrast, mutations that reduce the packing density at this interface such as small or flexible substitutions at L138 lead to LOF Orai1 channels. These phenotypes point to a model wherein the L138-T92 nexus acts as a lever/pivot point at the TM2-TM1 interface to mediate channel activation. Unusually, constitutively active currents arising from L138F and T92W mutations show fast CDI with kinetics comparable to WT Orai1 channels gated by STIM1. CDI of T92W Orai1 shows increased Ca^2+^ sensitivity and is not buffered by BAPTA, a chelator that prevents CDI of WT Orai1 channels. However, STIM1 co-expression normalizes the Ca^2+^ sensitivity of T92W Orai1 to that of WT channels gated by STIM1. These results have important implications for the CDI mechanism and the role of STIM1 in the feedback inhibition process.

### The T92-L138 motif regulates Orai1 activation

The finding that the L138-T92 motif regulates Orai1 gating is notable as this locus is located very closely to the H134 residue that functions as a steric “brake” at the interface between helices TM1, TM2, and TM3 to control Orai gating (Frischauf *et al*., 2017; Yeung *et al*., 2018). However, the dependence of sid-chain size substitutions at T92-L138 on Orai1 channel activity follows an inverse pattern of what is observed at H134 located one helical turn above this motif on TM2 **(*Figure 9A*)**. Introduction of bulky amino acids at T92 or L138 yield GOF channels, whereas small or flexible residues lead to LOF channels. At H134, by contrast, exactly the opposite is true: at this position small or flexible substitutions cause constitutive activity and large amino acids impede STIM1-mediated gating (Yeung *et al*., 2018).

**Figure 9.**
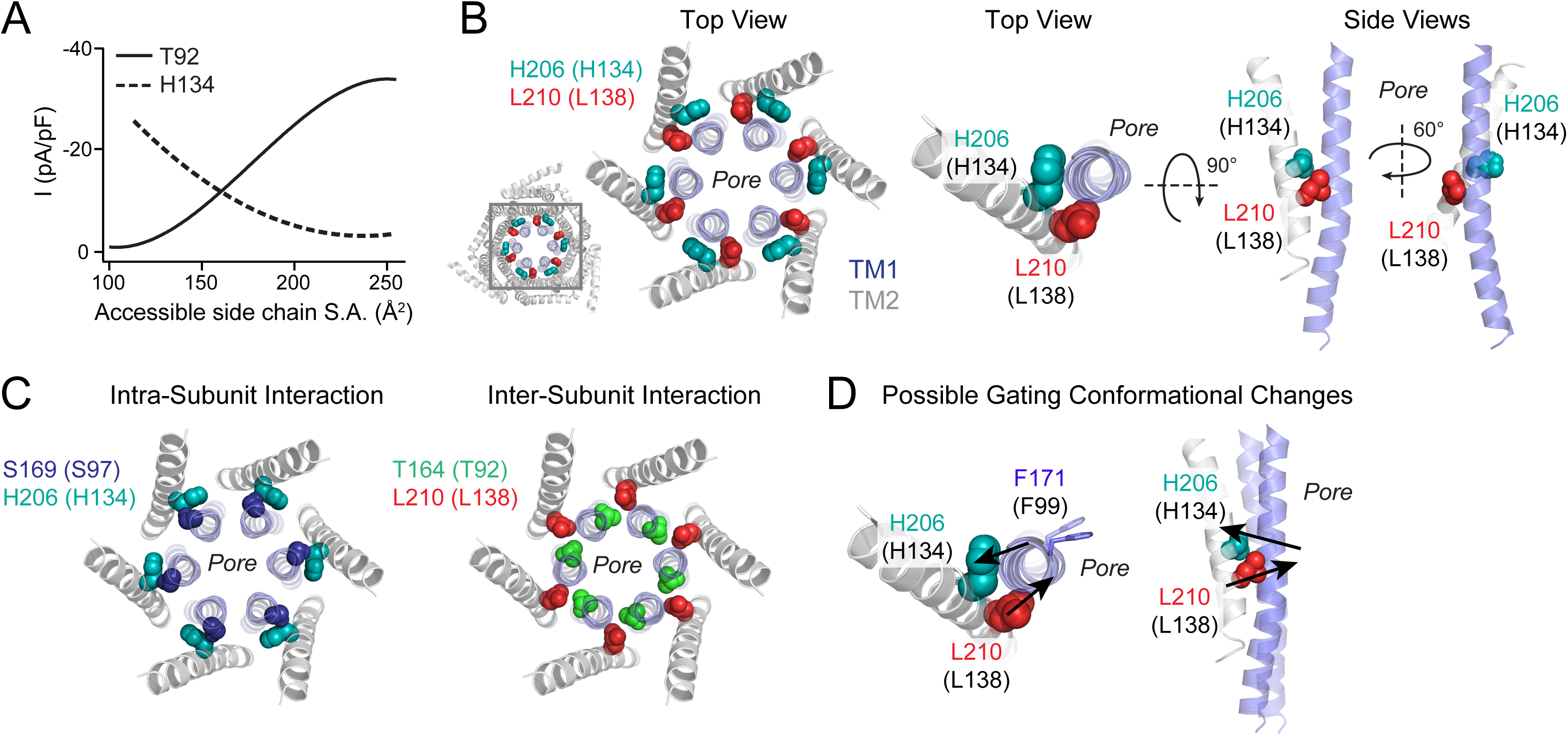
Schematic of the “brace” formed by the H134 and L138 residues around TM1. **(*A*)** Polynomial fits of T92 and H134 mutant channel activity plotted against side chain surface area shows opposite dependence of these two positions on size. For both fits, charged residues (D/E/K/R) and G/P were excluded from this analysis because of their propensity to break α-helical secondary structure. **(B)** Left: Top view of the dOrai hexameric channel with the relative positions at the TM1-TM2/3 ring interface with H134 (teal) and L138 (red) shown. Middle and right: Top and side views of residues H134 (teal spheres) and L138 (red spheres) at the TM1 (blue) and TM2 (grey) interface. The two amino acids are on either side of the pore helix, approximately one turn apart. One TM2 and TM1 helix from each subunit are displayed for simplicity. **(C)** Intra-subunit interaction of S97 (navy) with H134 (teal) and inter-subunit interaction of T92 (green) with L138 (red) are shown to highlight the different interfaces of these pairs of interactions. **(D)** Proposed gating conformational change of L138 and H134 associated with channel opening. H134 and L138 are positioned one helical turn away from each other on opposite sides of the TM1 pore helix. Amino acid substitutions at L138 that cause TM1 to be pushed inwards, or H134 to be drawn outwards, cause channel activation via TM1 pore helix twisting and pore dilation to open the channel gate.

What insight do the phenotypes of these mutants offer about the Orai1 channel activation process? The opposite dependencies of channel activity on side chain size at H134 and L138 implies that these two residues are complementary in their effects. Both residues are on TM2, which is tilted diagonally relative to the membrane and TM1 pore helix. H134 and L138 sit squarely at the crossover point between the two helices, and when viewed from the top, their side chains are in contact with opposite sides of TM1 **(*Figure 9B*)**. We postulate that their complementary functional role is related to the several key differences in their positioning – *i)* H134 and L138 are on opposite sides of TM1, *ii)* L138 is located one turn below H134, and *iii)* the S97-H134 interaction is intra-subunit (Yeung *et al*., 2018) while T92-L138 interaction is inter-subunit **(*Figure 9B,C*)**. We suggest that H134 and L138 function as pivot points or a “brace” for stabilizing the pore helix. In this scenario, introduction of steric pressure on TM1 (“pushing”) at L138 large amino acids substitutions or creation of a steric void (“pulling”) at H134 outward through small substitutions both lead to channel activation, while “pulling” at L138 or “pushing” at H134 abolishes channel function. Although speculative, this pattern also raises the possibility that during STIM1-mediated gating, the TM1 helix is pushed by the lower L138 and tilts towards the upper H134, leading to pore dilation in the hydrophobic stretch of the pore encompassing F99 that functions as the channel gate (Yamashita *et al*., 2017) **(*Figure 9D*)**. Moreover, because L138 and H134 are on different sides of TM1, they are also suitably positioned to help mediate rotation of the pore helix that accompanies pore dilation and channel opening (Yamashita *et al*., 2017; Yeung *et al*., 2018) **(*Figure 9D*)**.

### The T92-L138 motif regulates fast CDI: Implications for the CDI mechanism

Fast CDI is a distinguishing feature of CRAC channels and was first described in early recordings of CRAC currents in mast cells and T cells (Hoth & Penner, 1993; Zweifach & Lewis, 1995a). Although several reports have described effects of mutations in different regions of the CRAC channel on CDI, a broader molecular understanding of fast CDI remains elusive. Fundamental aspects of the process that are unclear include the identity of the Ca^2+^ sensor for CDI, the identity and nature of the inactivation gate that closes the pore, and the coupling mechanism between the Ca^2+^ sensor and the inactivation gate. A potential role for calmodulin (Mullins *et al*., 2009) or STIM1 (Derler *et al*., 2009; Scrimgeour *et al*., 2009; Mullins & Lewis, 2016) as the Ca^2+^ sensors mediating CDI were described in earlier studies, but subsequent evidence raised questions about the role of these molecules in sensing Ca^2+^ (Mullins & Lewis, 2016; Mullins *et al*., 2016), leaving the identity of the Ca^2+^ sensor a mystery. Nevertheless, a general consensus that STIM1 is necessary for driving CDI has emerged, and in particular, the ID region of STIM1 (encompassing residues 474-491) containing several acidic residues has been implicated as an essential domain for CDI. STIM1 mutants lacking ID_STIM1_ fail to show CDI, and mutating specific residues within the region accelerate or in some cases diminish CDI (Mullins & Lewis, 2016; Mullins *et al*., 2016). Within Orai1, mutational analysis and domain swapping experiments between different Orai isoforms have implicated the cytosolic domains, in particular the Orai1 N- and the C-termini (Lee *et al*., 2009; Srikanth *et al*., 2010; Frischauf *et al*., 2011; Mullins *et al*., 2016). The most generally accepted view is that the inactivation domain of STIM1 is allosterically coupled to the inner pore to mediate CDI (Mullins & Lewis, 2016; Mullins *et al*., 2016). In particular, the aromatic residues Y80 and W76 and to a lesser extent, the three positively charged residues (R91, K87, and R83), have been strongly implicated (Mullins *et al*., 2016) raising the possibility that this region of the inner pore may function as the channel gate for CDI.

Against this backdrop, the finding that Orai1 L138F channels and T92W channels exhibit rapid Ca^2+^-dependent inactivation during hyperpolarizing steps in the absence of STIM1 provide several new insights on the mechanism of the inactivation process. T92W Orai1 inactivation shares several features with rapid CDI of CRAC channels. These include dependence on extracellular divalent cations, increasing inactivation driven by increasing extracellular Ca^2+^ concentrations, and hyperpolarizing pulses that increase the driving force for Ca^2+^ entry. Moreover, the kinetics of inactivation of T92W and L138F Orai1 channels occurs with a biexponential time course similar to that observed in native CRAC channels in EGTA (ρ_fast_ and ρ_slow_ of ∼10 and 130 ms respectively). Additionally, the E106D Orai1 mutation, which abrogates CDI of WT Orai1 channels gated by STIM1 (Yamashita *et al*., 2007), also eliminates CDI of T92W Orai1. These similarities indicate that the rapid inactivation of T92W and L138F shares a similar gating mechanism as CDI in native CRAC channels.

The finding that T92W and L138F Orai1 variants exhibit fast CDI in the absence of STIM1 have several important implications for the mechanism of CDI. First, this result indicates that STIM1 is not essential for mediating CDI, nor does it function as the Ca^2+^ sensor. Rather, the results suggest that the Ca^2+^ sensor is likely located within the Orai1 protein itself, or a closely associated accessory subunit. We favor the idea that the Ca^2+^ binding site is located on Orai1 itself based on previous observations in Orai2 and Orai3 (Lee *et al*., 2009) reaffirmed here for Orai1, indicating that the Orai1 C-terminus has a key role in mediating CDI.

A surprising finding is that CDI of T92W Orai1 is much more prominent in BAPTA-containing internal solutions compared to EGTA-based solutions. This is the exact reverse of what is seen for WT Orai1 channels. Our analysis suggests that this is due to enhanced Ca^2+^ sensitivity of T92W Orai1 channels for CDI compared to WT Orai1 channels. Remarkably, co-expression of STIM1 with T92W Orai1 channels “normalizes” the EGTA/BAPTA dependence of inactivation of the mutant channels reminiscent of the normalization of ion selectivity and permeation of mutant V102C/A Orai1 channels by STIM1 (McNally *et al*., 2012). This effect appears to be mediated via decrease in the Ca^2+^ sensitivity of the CDI, indicating that the role of STIM1 for CDI is to modulate the Ca^2+^-sensitivity of the inactivation process. A change in Ca^2+^ sensitivity of CDI could presumably occur via STIM1-driven change in the conformation of the domain containing the Ca^2+^ binding site at the Orai1 C-terminus, which our results indicate is a key region that mediates T92W Orai1 CDI. The increased Ca^2+^ sensitivity of T92W and L138F CDI likely also explains the unusually high I_Na+_/I_Ca2+_ current ratios observed for these mutants as inactivation is relieved in Na^+^-containing solutions. Moreover, CDI of L138F may also explain previous observations indicating that this mutation raises resting cytoplasmic [Ca^2+^] to a lesser degree than other gain-of-function human mutations (e.g. S97C) and evokes tubular aggregate myopathy but not hypocalcemia (Endo *et al*., 2015).

The Orai1 C-terminus harbors several acidic residues in the close vicinity of L273 and L276 which are critical for STIM1 binding. Interaction of STIM1 with the Orai1 C-terminus would naturally be expected to alter the conformation of the Orai1 C-terminus, and therefore induce the change in Ca^2+^ sensitivity of CDI of T92W Orai1 in response to STIM1 binding. Additionally, we find that mutations in the Orai1 N-terminus (W76E, Y80E, R83E) that were previously found to abrogate CDI in the presence of STIM1 (Mullins & Lewis, 2016; Mullins *et al*., 2016), substantially accelerated and increased CDI of T92W Orai1 channels. These phenotypes reinforce that notion that the inner pore is a key molecular determinant of CDI in Orai1 channels, but the acceleration of clearly CDI indicates that this region is unlikely to function as the inactivation gate. Together, these lines of evidence indicate that CDI of Orai1 occurs independently of STIM1 with the Ca^2+^ binding site likely located within Orai1 itself, possibly at the Orai1 C-terminus. The Orai1 C-terminus contains a cluster of acidic residues which could function as a distributed Ca^2+^ sensing region, and consistent with this idea, previous evidence has implicated some acidic residues for CDI of Orai2 channels (Lee *et al*., 2009). Since the primary STIM1 binding site in Orai1 is located within this same region in the C-terminus (Park *et al*., 2009), STIM1 binding would naturally be expected to tune the Ca^2+^ sensitivity of the CDI process as explicitly shown in our results. The precise identity and nature of the Ca^2+^ binding site and how Ca^2+^ binding to the sensor is communicated to the pore to evoke channel closure remain to be understood but the results here provide a way forward to address these questions in a simplified one-component system using a constitutively active Orai1 mutant that shows CDI in the absence of STIM1.

## Materials and Methods

### Key Resources Table

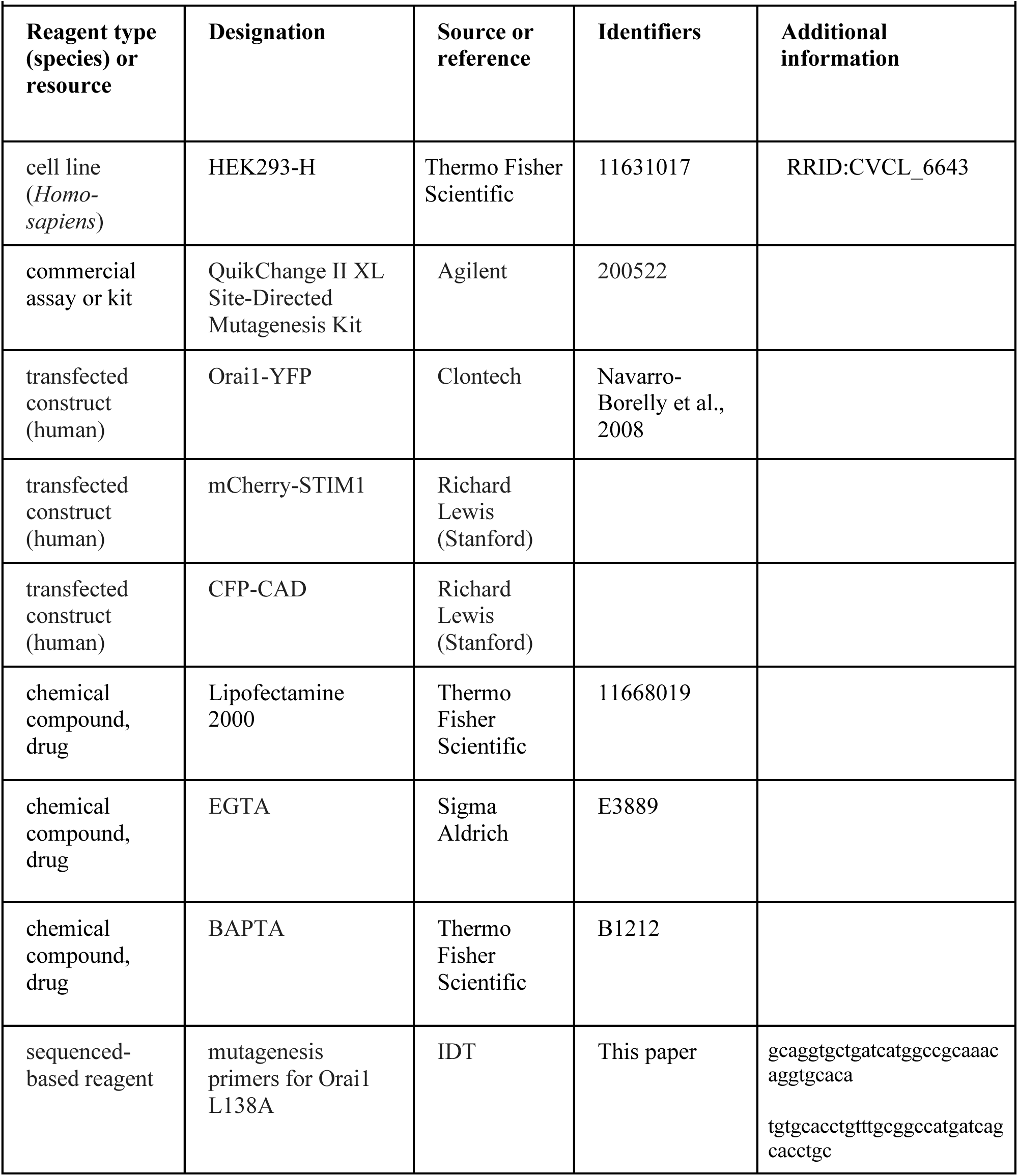
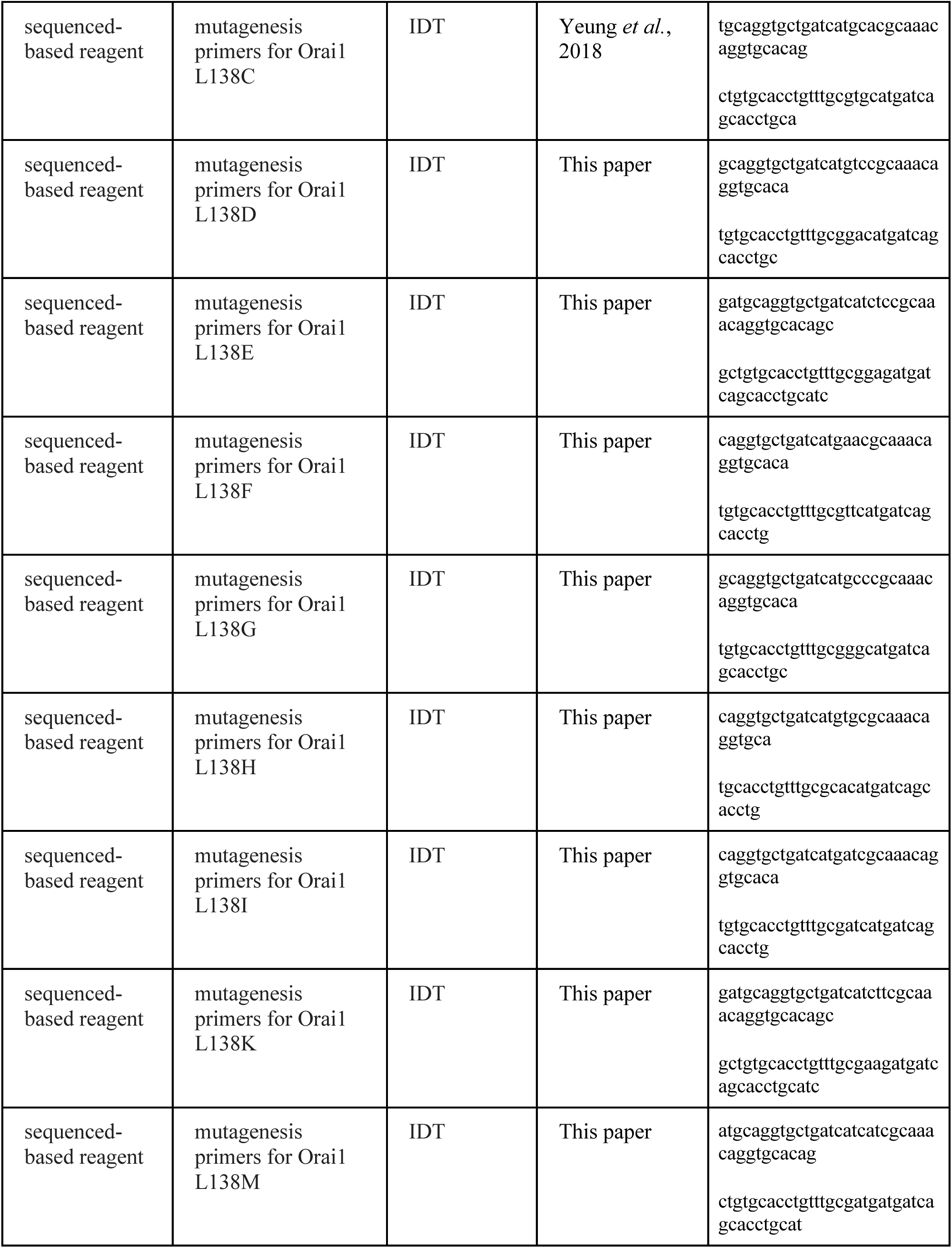
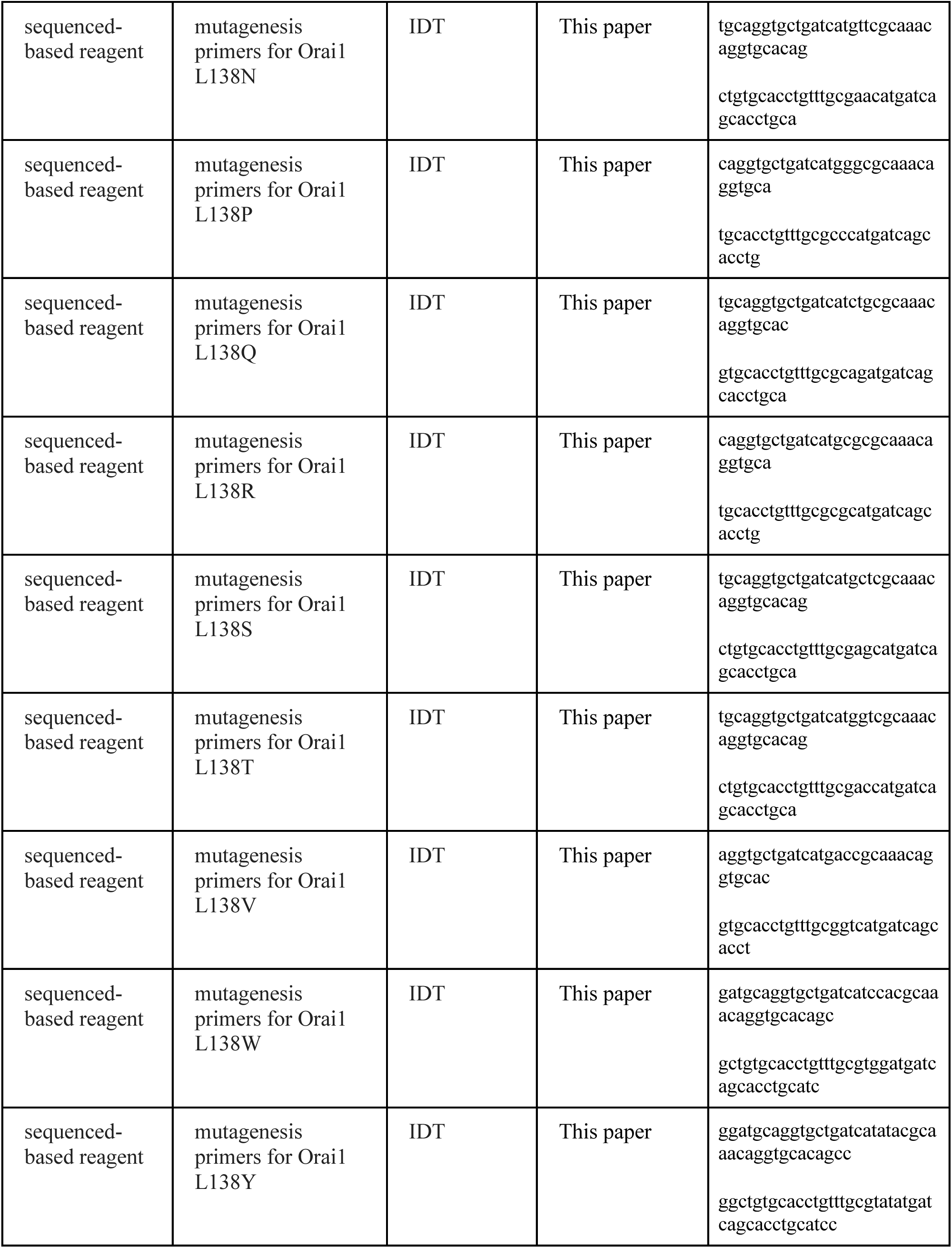
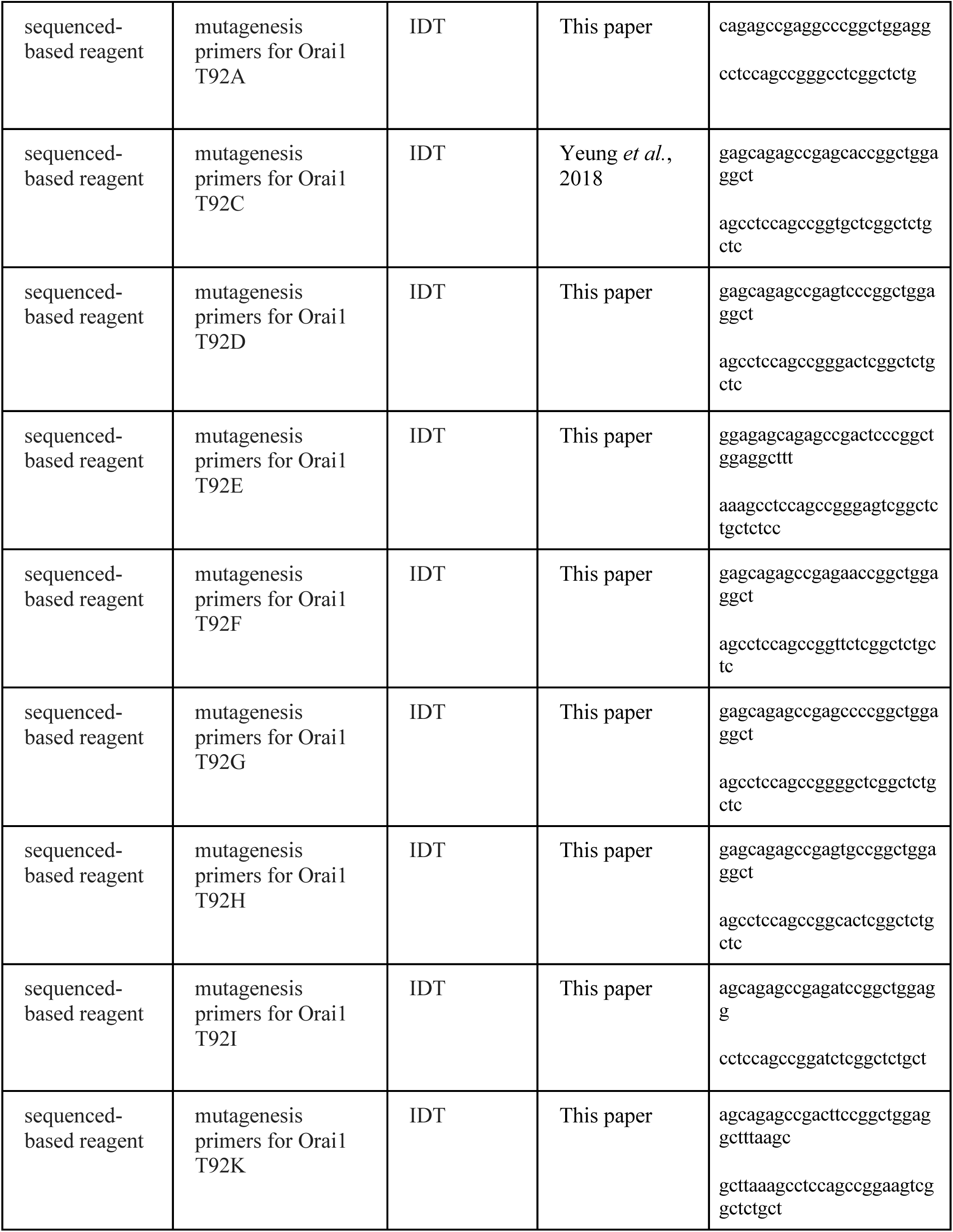
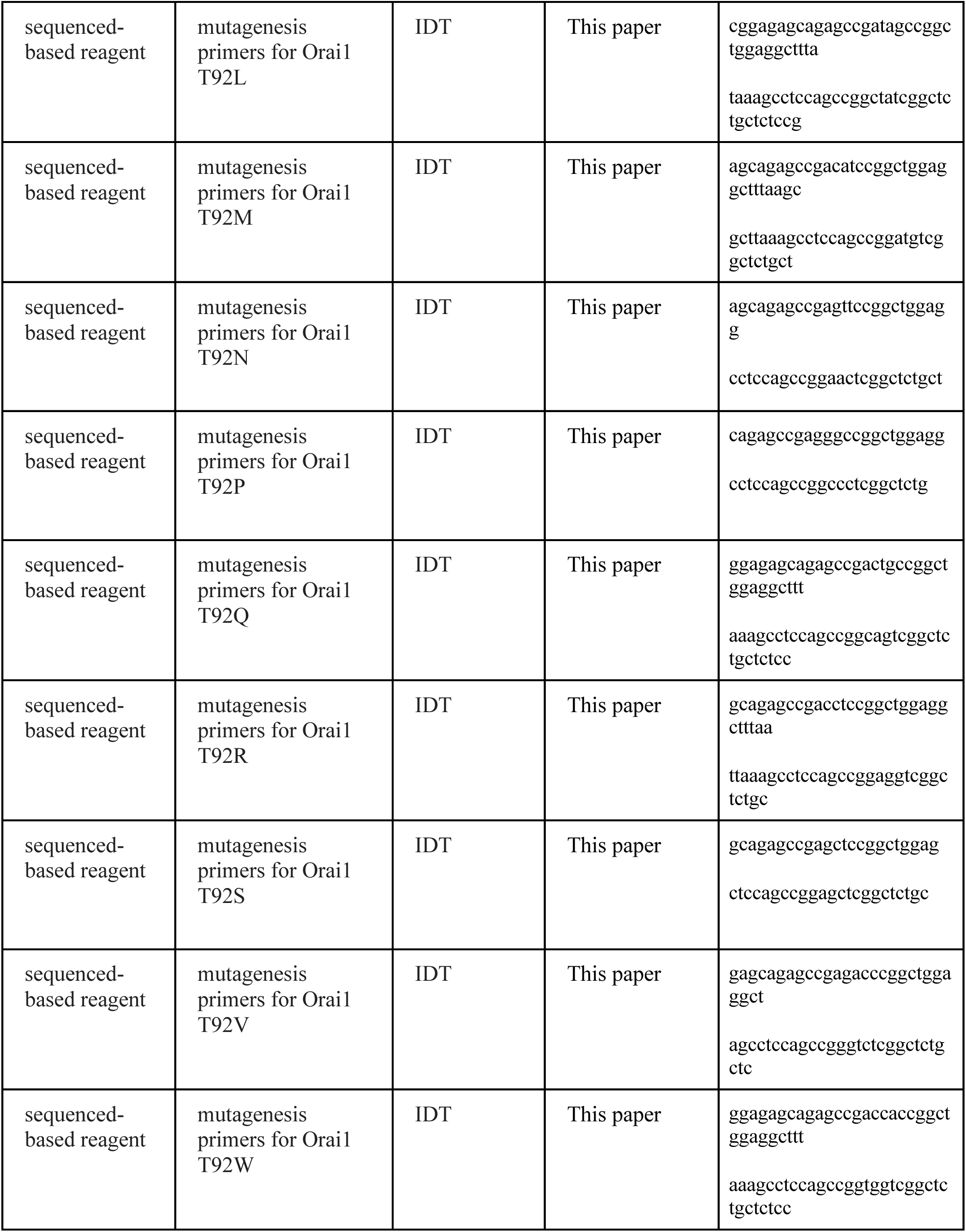
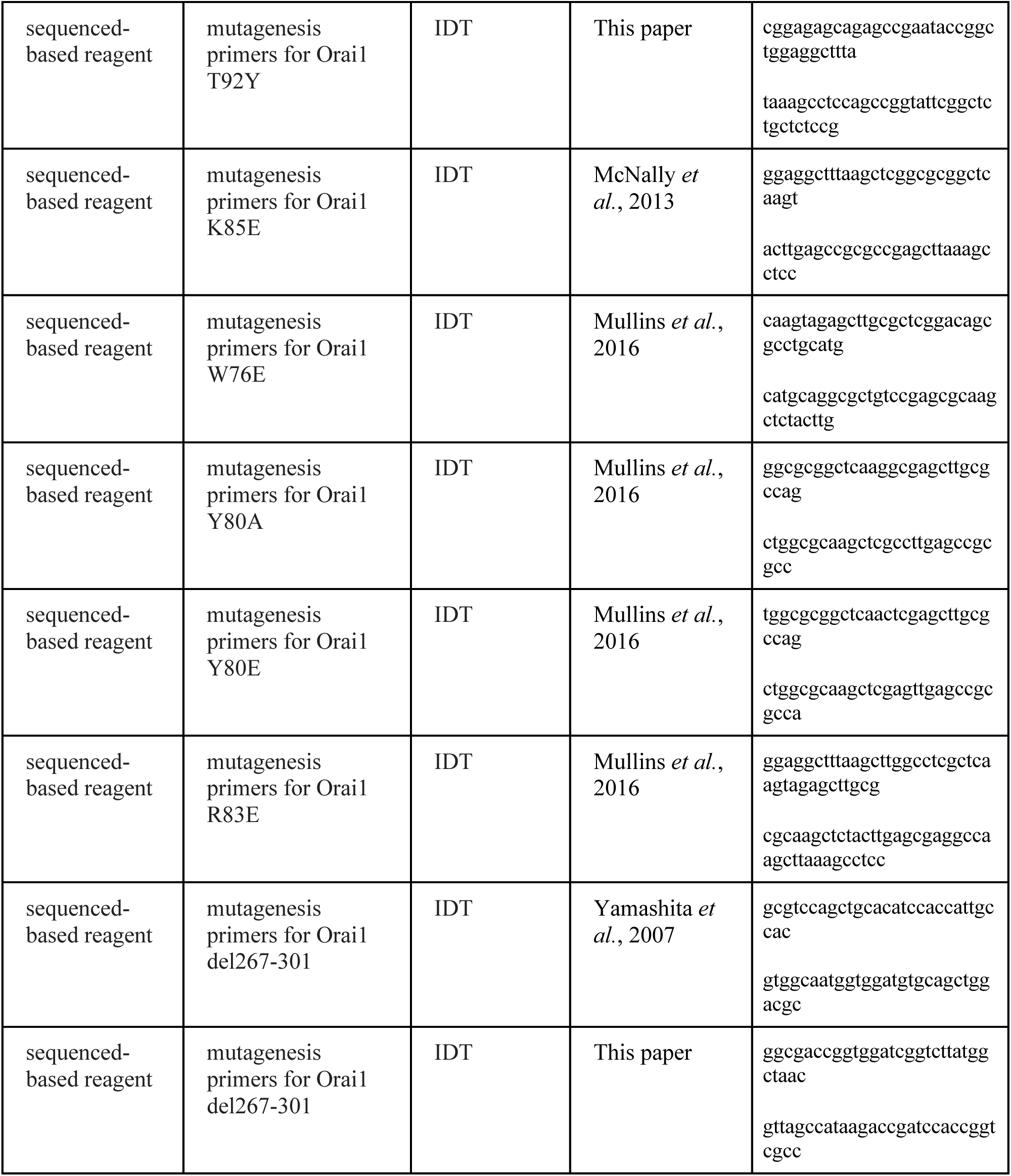

### Cells

HEK293-H cells were maintained in suspension at 37°C with 5% CO_2_ in CD293 medium supplemented with 4 mM GlutaMAX (Invitrogen). For imaging and electrophysiology, cells were plated onto poly-L-lysine coated coverslips one day before transfection and grown in a medium containing 44% DMEM (Corning), 44% Ham’s F12 (Corning), 10% fetal bovine serum (HyClone), 2 mM glutamine, 50 U/ml penicillin and 50 µg/ml streptomycin.

### Plasmids and transfections

The Orai1 mutants employed for electrophysiology were engineered into a pEYFP-N1 vector (Clonetech) to produce C-terminally tagged Orai1-YFP proteins (Navarro-Borelly *et al*., 2008). mCherry-STIM1 and CFP-CAD were kind gifts of Dr. R. Lewis (Stanford University, USA). All mutants were generated by the QuikChange Mutagenesis Kit (Agilent Technologies) and the mutations were confirmed by DNA sequencing. For electrophysiology, the indicated Orai1 constructs were transfected into HEK293-H cells either alone (200 ng DNA per coverslip) or together with STIM1 (100 ng Orai1 and 500 ng STIM1 DNA per coverslip). For FRET microscopy experiments, cells were transfected with Orai1-YFP and CFP-CAD constructs (100 ng each per coverslip). All transfections were performed using Lipofectamine 2000 (Thermo Fisher Scientific) 24-48 hours prior to electrophysiology or imaging experiments.

### Solutions and Chemicals

The standard 20 mM Ca^2+^ extracellular Ringer’s solution used for electrophysiological experiments contained 135 mM NaCl, 4.5 mM KCl, 20 mM CaCl_2_, 1 mM MgCl_2_, 10 mM D-glucose, and 5 mM HEPES (pH 7.4 with NaOH). 110 mM Ca^2+^ solution contained 110 mM CaCl_2_, 10 mM D-glucose, and 5 mM HEPES (pH 7.4 with NaOH). The divalent-free (DVF) solution contained 150 mM NaCl, 10 mM HEDTA, 1 mM EDTA, and 10 mM HEPES (pH 7.4 with NaOH). 10 mM TEA-Cl was added to prevent contamination from voltage-gated K^+^ channels. All internal solutions contained 8 mM MgCl_2_ and 10 mM HEPES (pH 7.2 with CsOH). The standard 8 mM BAPTA internal solution contained 135 mM Cs aspartate and 8 mM BAPTA, and the 0.8 mM BAPTA solution contained 145 mM Cs aspartate and 0.8 BAPTA. The 10 mM EGTA solution contained 130 mM Cs aspartate and 10 mM EGTA, and the 20 mM EGTA solution contained 110 mM Cs aspartate and 20 mM EGTA.

### Electrophysiology

Currents were recorded in the standard whole-cell configuration at room temperature on an Axopatch 200B amplifier (Molecular Devices) interfaced to an ITC-18 input/output board (Instrutech). Routines developed by R. S. Lewis (Stanford) on the Igor Pro software (Wavemetrics) were employed for stimulation, data acquisition and analysis. Data are corrected for the liquid junction potential of the pipette solution relative to Ringer’s in the bath (−10 mV). The holding potential was +30 mV. The standard voltage stimulus consisted of a 100- ms step to –100 mV followed by a 100-ms ramp from –100 to +100 mV applied at 1 s intervals. For voltage families, steps to −120 mV, − 100 mV, −80 mV, and −60 mV were 300 ms each. In the paired-pulse experiment, the holding potential was +30 mV and the two steps were to −100 mV for 300 ms each separated by a step to +100 mV for 200 ms in between the two hyperpolarizing steps. In experiments where Orai1 was co-expressed with STIM1, I_CRAC_ was typically activated by passive depletion of ER Ca^2+^ stores by intracellular dialysis of 8 mM BAPTA. All currents were acquired at 5 kHz and low pass filtered with a 1 kHz Bessel filter built into the amplifier. All data were corrected for leak currents collected in 100-200 µM LaCl_3_.

### Data analysis

Analysis of current amplitudes was typically performed by measuring the peak currents during the −100 mV pulse. Specific mutants were categorized as gain-of-function if their currents exceeded 2 pA/pF, which is more than ten times the current density of WT Orai1 without STIM1. Reversal potentials were measured from the average of several leak-subtracted sweeps in each cell. For CDI, the extent of inactivation was determined from the relative decrease in current (relative to the peak current) during the voltage pulse and quantified as (1−*I_ss_*/*I_peak_*) where Iss is the current at the end of the 300 ms hyperpolarizing step and Ipeak is the peak current immediately following the hyperpolarizing step. The time course of CDI was fit with a double-exponential function and the fast and slow time constants (𝝉_fast_ and 𝝉_slow_) were determined from the fits. All fitting was done using the built-in routines in Igor Pro v6.12. All data are expressed as means ± SEM. For datasets with two groups, statistical analysis was performed with two-tailed t test to compare between control and test conditions. For datasets with greater than two groups, one-way ANOVA followed by Tukey post-hoc test was used to compare groups. Statistical analysis was performed with a confidence level of 95%, and results with P < 0.05 were considered statistically significant. Significance is denoted as *p<0.05, **p<0.01, ***p<0.001.

### Atomic packing analysis

Atomic packing analysis was performed as in our previous study (Yeung *et al*., 2018). Briefly, it carried out using the programs REDUCE and PROBE that simulates rolling a 0.25Å radius sphere along the van der Waals surfaces. Locations where the probe sphere contacts two surfaces are marked (with a ‘dot’) that classifies whether the surfaces are in wide contact, close contact, overlapped, or clashing. The resulting contact dot scores were summed for all atoms of each residue and displayed using PyMOL on a heat map that shows the degree of contacts.

### FRET microscopy

HEK293-H cells transfected with Orai1-YFP and CFP-CAD DNA constructs were imaged using wide-field epifluorescence microscopy on an IX71 inverted microscope (Olympus, Center Valley, PA). Cells were imaged with a 60X oil immersion objective (UPlanApo NA 1.40), a 175 W Xenon arc lamp (Sutter, Novatao, CA), and excitation and emission filter wheels (Sutter, Novato, CA). At each time point, three sets of images (CFP, YFP, and FRET) were captured on a cooled EM-CCD camera (Hamamatsu, Bridgewater, NJ) using optical filters specific for the three images as previously described. Image acquisition and analysis was performed with SlideBook software (Imaging Innovations Inc., Denver, CO). Images were captured at exposures of 100-500 ms with 1X1 binning. Lamp output was attenuated to 25% by a 0.6 ND filter in the light path to minimize photobleaching. All experiments were performed at room temperature.

FRET analysis was performed as previously described (Navarro-Borelly *et al*., 2008). The microscope-specific bleed-through constants (a=0.12; b=0.008; c=0.002 and d=0.33) were determined from cells expressing cytosolic CFP or YFP alone. The apparent FRET efficiency was calculated from background-subtracted images using the formalism (Zal & Gascoigne, 2004):

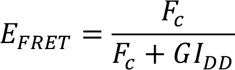

where *F_c_ = I_DA_ - aI_AA_ - dI_DD_*

*Idd*, *Iaa* and *Ida* refer to the background subtracted CFP, YFP, and FRET images, respectively. The instrument dependent *G* factor had the value 1.85 ± 0.1. E-FRET analysis was restricted to cells with YFP/CFP ratios in the range of 2-6 to ensure that E-FRET was compared across identical acceptor to donor ratios, and measurements were restricted to regions of interest drawn at the plasma membrane.

## Acknowledgements

We thank members of the laboratory for helpful discussions. This work was supported by NIH grants R01 NS057499 to M.P. P.S.-W.Y. was supported by NIH predoctoral fellowship F31NS101830.

## Author contributions

P.S.-W.Y. and M.Y. generated and functionally characterized the Orai1 mutants by patch-clamp electrophysiology, atomic packing analysis, and FRET imaging. M.P. supervised the work, analyzed data, and wrote the manuscript. All authors were contributed to the experimental design, data analysis, and writing of the paper.

## Competing financial interests

The authors declare that we have no competing financial interests.

---

**Figure 1 – Figure Supplement 1.**
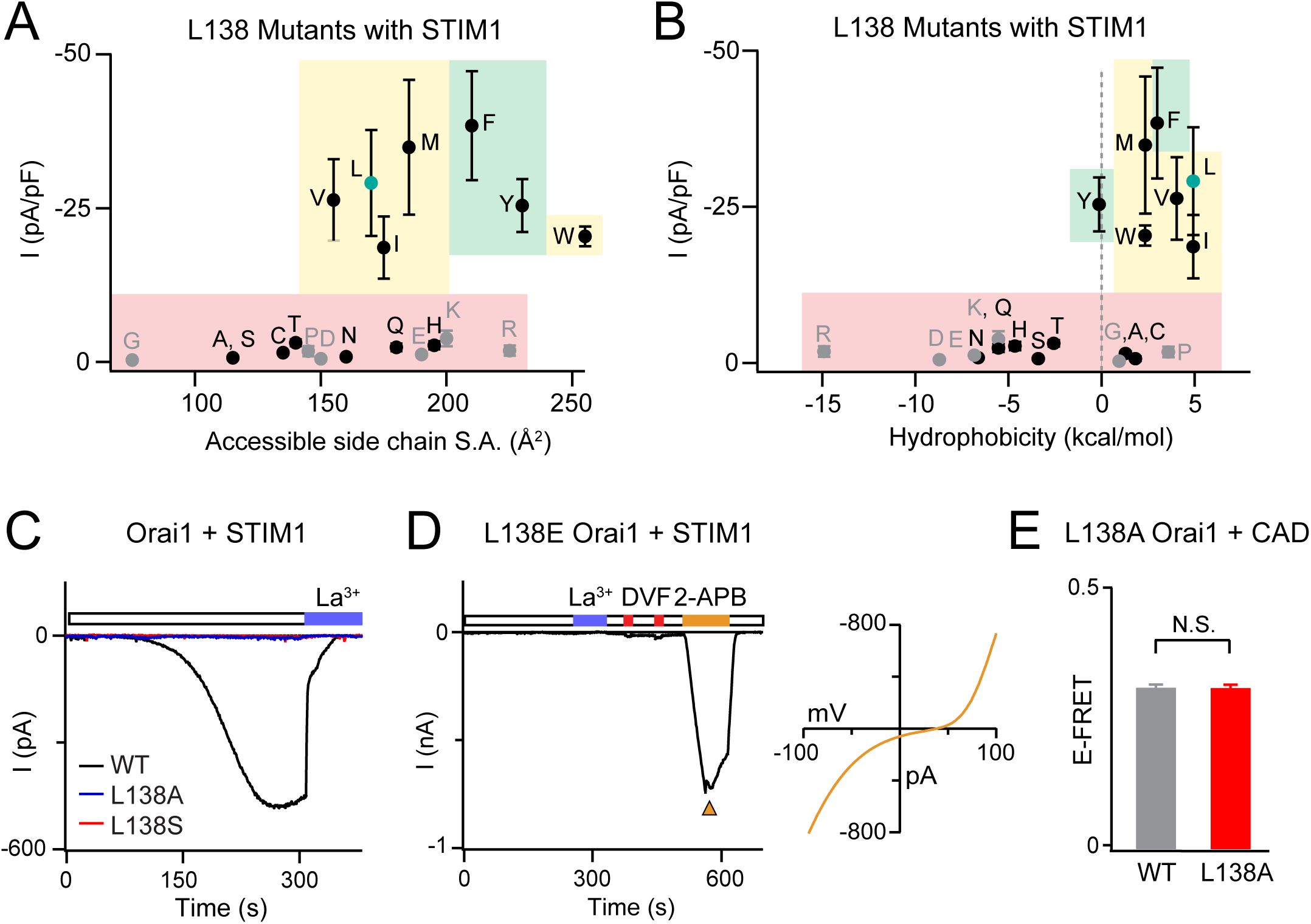
Analysis of L138 Orai1 currents in the presence of STIM1 co-expression. **(A-B)** Current densities of L138 mutants expressed with STIM1 are plotted against size chain size and hydrophobicity. Small and flexible substitutions produce LOF channels. Green, yellow, and red shading highlights GOF mutants, store-operated, and LOF mutants, respectively. N = 4-7 cells. Values are mean ± S.E.M. **(C)** Unlike WT Orai1, L138A and L138S mutants do not show current induction over time with STIM1 co-expression. **(D)** L138E mutant cannot be gated by STIM1 but is strongly activated by Orai1 modulator 2-APB (50 μM). **(E)** There is no defect in L138A Orai1 interaction with CAD as measured by E-FRET between L138A Orai1-YFP and CFP-CAD, indicating that loss of gating in the L138A mutant is not due to lack of membrane expression or defect in interaction with STIM1. N = 51-57 cells. Values are mean ± S.E.M.

**Figure 2 – Figure Supplement 1.**
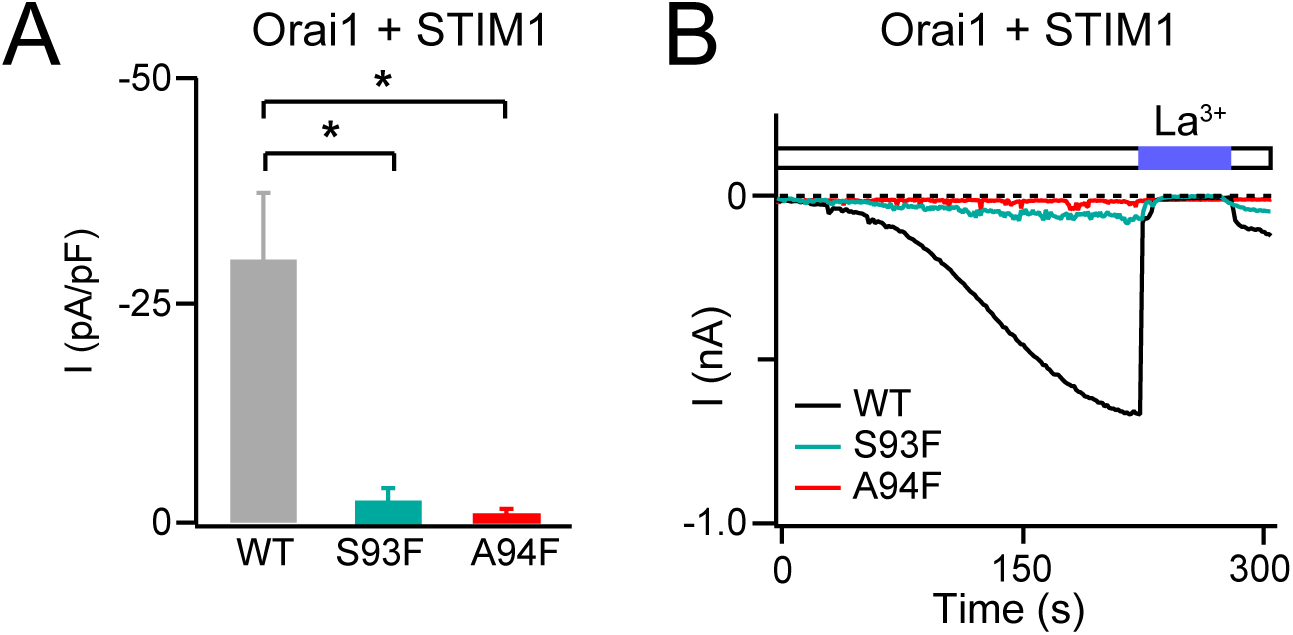
S93F and A94F mutations impede channel activation by STIM1. **(A-B)** Average current densities of S93F and A94F Orai1 channels expressed with STIM1. S93F and A94F Orai1 are non-functional even when co-expressed with STIM1. N = 4 cells. Values are mean ± S.E.M. *: p<0.05 by t-test

**Figure 3 – Figure Supplement 1.**
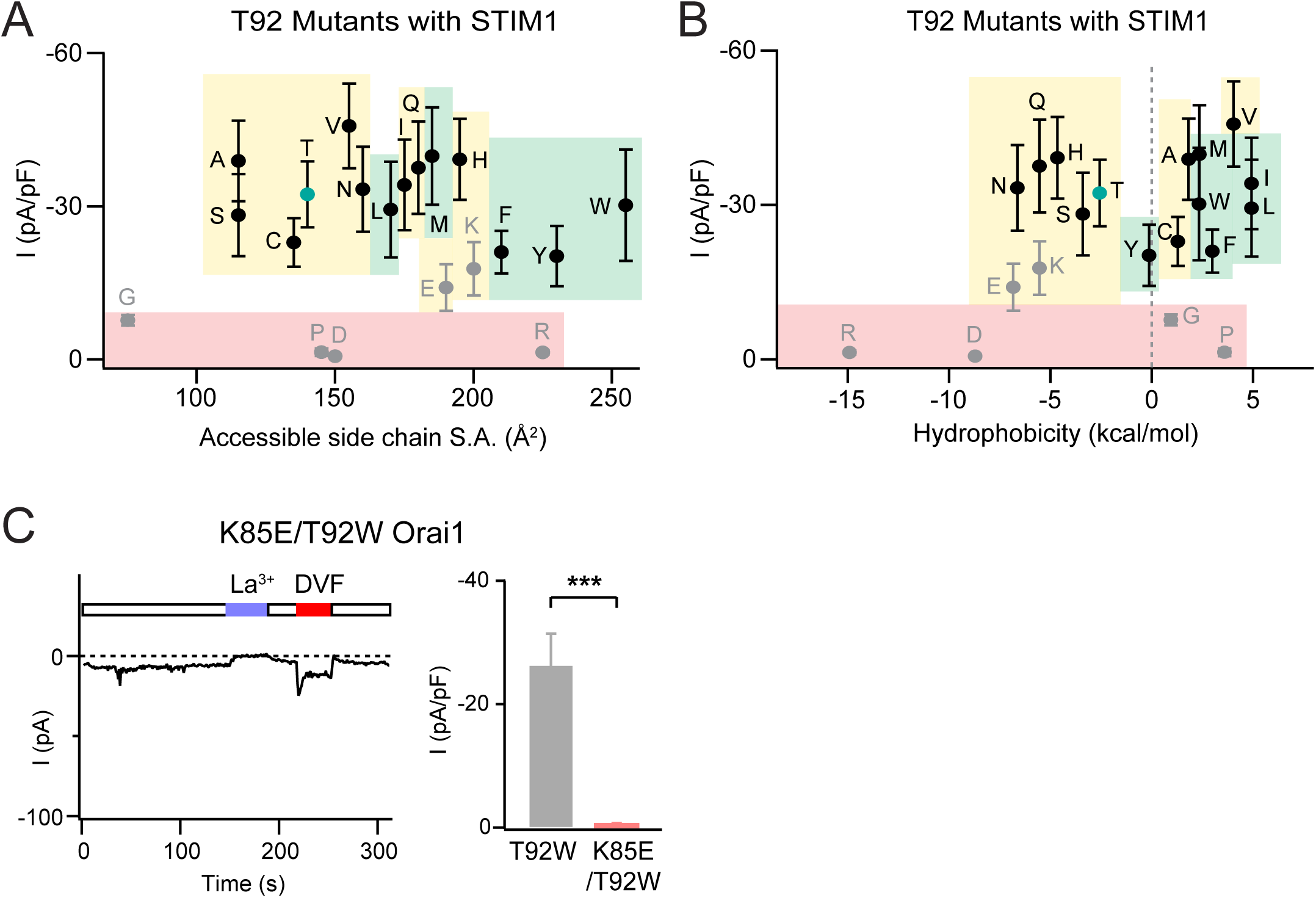
Analysis of T92 Orai1 mutant currents in the presence of STIM1. **(A-B)** The current densities of T92 mutants with STIM1 co-expression plotted against size-chain size and hydrophobicity. Green, yellow, and red shading highlights GOF mutants, store-operated, and LOF mutants, respectively. N = 4-8 cells. Values are mean ± S.E.M. **(C)** Like other known GOF mutants (Yeung *et al*., 2018), the current of T92W is abrogated by the addition of LOF N-terminal mutation K85E. N =5-8 cells. Values are mean ± S.E.M. ***: p<0.001 by t-test.

**Figure 4 – Figure Supplement 1.**
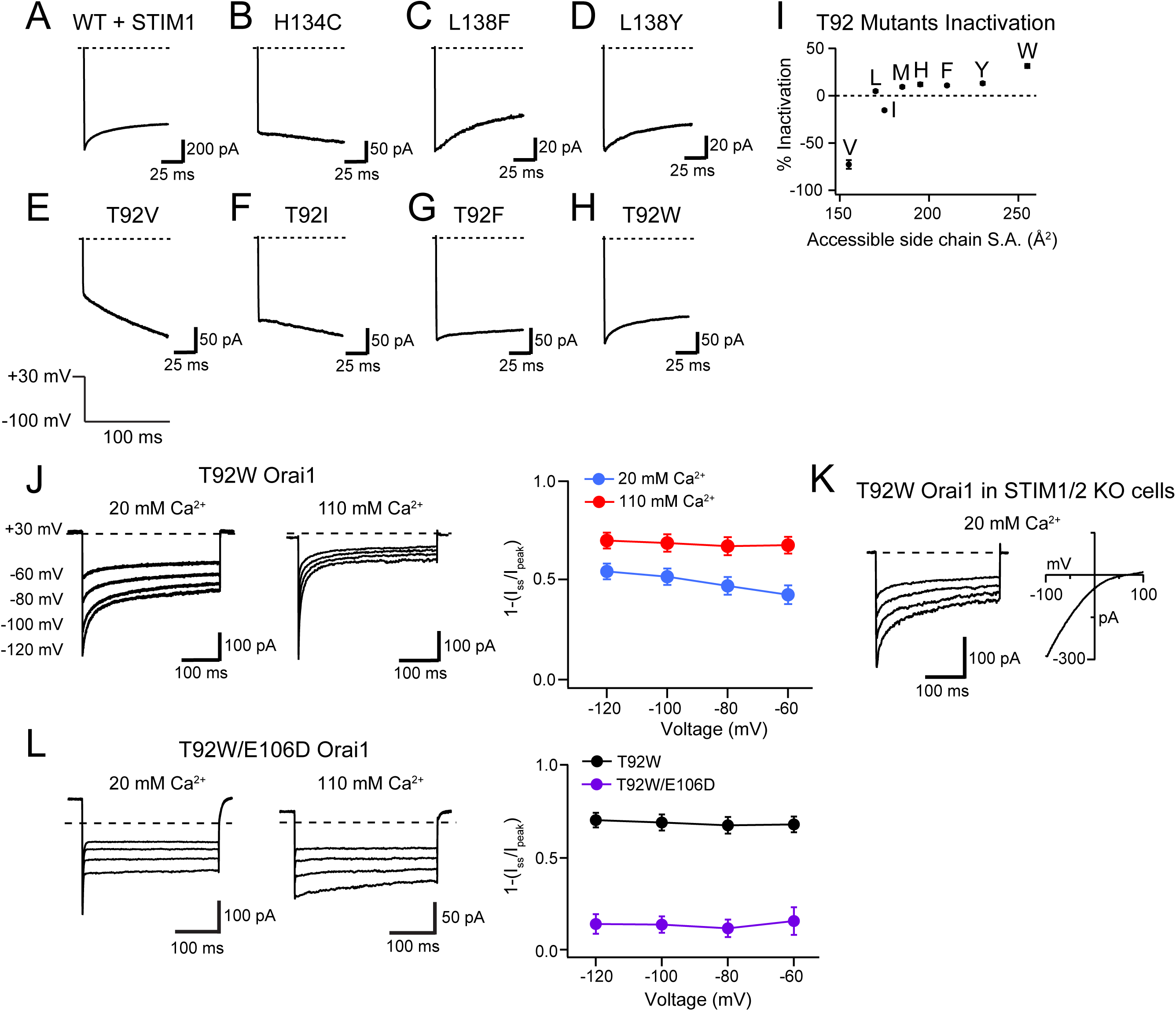
Analysis of inactivation of T92 mutants. **(A-H)** Example traces showing inactivation seen in STIM1-gated WT and L138 or T92 mutant channels during a 100 ms hyperpolarizing step from +30 mV to −100 mV. **(B)** GOF mutants such as H134C typically do not inactivate without STIM1. **(C-D)** L138F and L138Y mutants show inactivation during the 100 ms hyperpolarizing step with kinetics comparable to CDI of WT Orai1 channels. **(E-I)** Inactivation of T92 mutants is positively correlated with the side-chain surface area (S.A.) at this position. T92V shows slight potentiation similar other previously described GOF mutants, whereas T92W inactivates 30% over the 100 ms hyperpolarization step to −100 mV with kinetics similar to STIM1-gated channels. **(J)** Inactivation of T92W mutant is enhanced by raising extracellular Ca^2+^ to 110 mM. **(K)** T92W Orai1 is constitutively active and displays inactivation in STIM1/2 double knock-out HEK cells, indicating that the activation and inactivation of this mutant is not dependent on STIM. **(L)** As previously shown for the E106D Orai1 single mutant gated by STIM1 (Yamashita et al., 2007), T92W/E106D Orai1 shows Ca^2+^-dependent block of Na^+^ current in 20 mM external Ca^2+^ and minimal inactivation in 110 mM Ca^2+^ solution. Inactivation of the T92W mutant in 110 mM Ca^2+^ solution with and without E106D. N = 5-17 cells. Values are mean ± S.E.M.

**Figure 8 – Figure Supplement 1.**
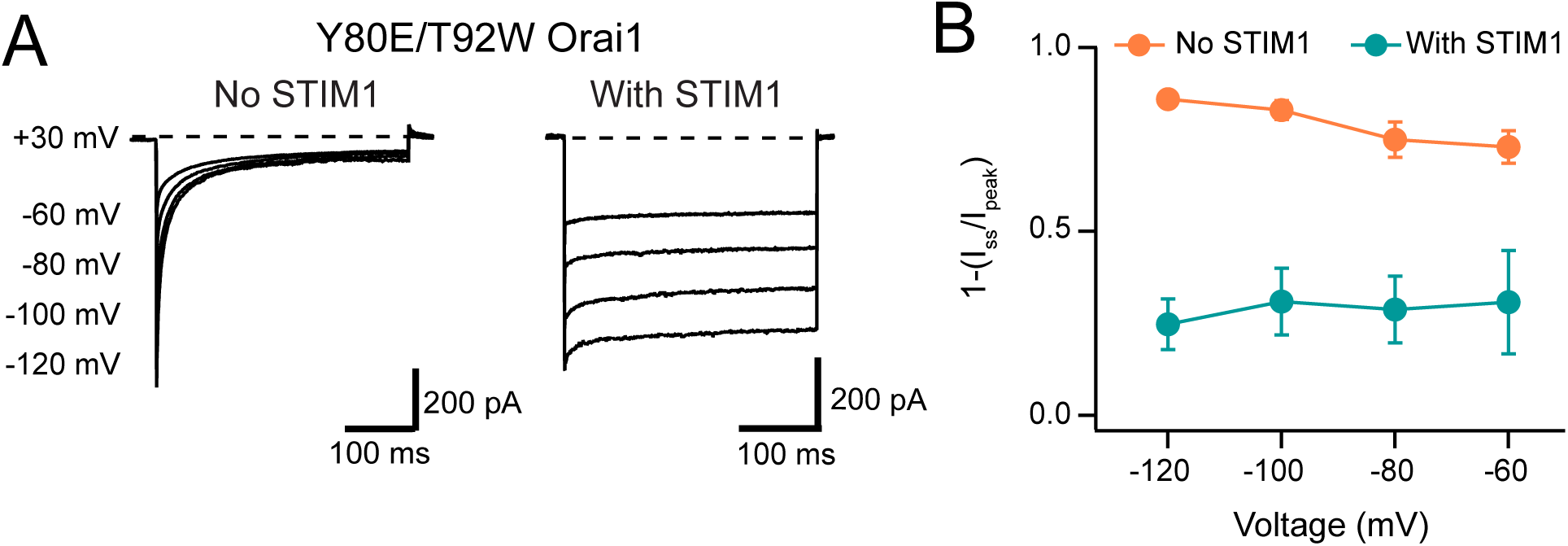
STIM1 modulates CDI of Y80E/T92W Orai1. **(A)** Example traces of Y80E/T92W voltage family steps without and with STIM1 co-expression. **(B)** Summary data of Y80E/T92W Orai1 inactivation modulation by STIM1. The example trace and quantification shown without STIM1 is the same data as in Fig. 7 E-F. N = 4 cells. Values are mean ± S.E.M.

## Notes

### Competing Interest Statement

The authors have declared no competing interest.

## References

Bergsmann J, Derler I, Muik M, Frischauf I, Fahrner M, Pollheimer P, Schwarzinger C, Gruber HJ, Groschner K & Romanin C. (2011). Molecular determinants within N terminus of Orai3 protein that control channel activation and gating. J Biol Chem 286, 31565–31575.

Bohm J, Bulla M, Urquhart JE, Malfatti E, Williams SG, O’Sullivan J, Szlauer A, Koch C, Baranello G, Mora M, Ripolone M, Violano R, Moggio M, Kingston H, Dawson T, DeGoede CG, Nixon J, Boland A, Deleuze JF, Romero N, Newman WG, Demaurex N & Laporte J. (2017). ORAI1 Mutations with Distinct Channel Gating Defects in Tubular Aggregate Myopathy. Human mutation 38, 426–438.

Bulla M, Gyimesi G, Kim JH, Bhardwaj R, Hediger MA, Frieden M & Demaurex N. (2019). ORAI1 channel gating and selectivity is differentially altered by natural mutations in the first or third transmembrane domain. J Physiol 597, 561–582.

Derler I, Butorac C, Krizova A, Stadlbauer M, Muik M, Fahrner M, Frischauf I & Romanin C. (2018). Authentic CRAC channel activity requires STIM1 and the conserved portion of the Orai N terminus. J Biol Chem 293, 1259–1270.

Derler I, Fahrner M, Muik M, Lackner B, Schindl R, Groschner K & Romanin C. (2009). A Ca^2+^ release-activated Ca^2+^ (CRAC) modulatory domain (CMD) within STIM1 mediates fast Ca^2+^ -dependent inactivation of ORAI1 channels. J Biol Chem 284, 24933–24938.

Emrich SM, Yoast RE, Xin P, Zhang X, Pathak T, Nwokonko R, Gueguinou MF, Subedi KP, Zhou Y, Ambudkar IS, Hempel N, Machaca K, Gill DL & Trebak M. (2019). Cross-talk between N-terminal and C-terminal domains in stromal interaction molecule 2 (STIM2) determines enhanced STIM2 sensitivity. J Biol Chem 294, 6318–6332.

Endo Y, Noguchi S, Hara Y, Hayashi YK, Motomura K, Miyatake S, Murakami N, Tanaka S, Yamashita S, Kizu R, Bamba M, Goto Y, Matsumoto N, Nonaka I & Nishino I. (2015). Dominant mutations in ORAI1 cause tubular aggregate myopathy with hypocalcemia via constitutive activation of store-operated Ca^2+^ channels. Hum Mol Genet 24, 637–648.

Feske S. (2010). CRAC channelopathies. Pflugers Archiv : European journal of physiology 460, 417–435.

Fierro L & Parekh AB. (1999a). The effects of interfering with GTP-binding proteins on the activation mechanism of calcium release-activated calcium current. Pflugers Arch 437, 547–552.

Fierro L & Parekh AB. (1999b). Fast calcium-dependent inactivation of calcium release-activated calcium current (CRAC) in RBL-1 cells. J Membr Biol 168, 9–17.

Frischauf I, Litvinukova M, Schober R, Zayats V, Svobodova B, Bonhenry D, Lunz V, Cappello S, Tociu L, Reha D, Stallinger A, Hochreiter A, Pammer T, Butorac C, Muik M, Groschner K, Bogeski I, Ettrich RH, Romanin C & Schindl R. (2017). Transmembrane helix connectivity in Orai1 controls two gates for calcium-dependent transcription. Sci Signal 10.

Frischauf I, Schindl R, Bergsmann J, Derler I, Fahrner M, Muik M, Fritsch R, Lackner B, Groschner K & Romanin C. (2011). Cooperativeness of Orai cytosolic domains tunes subtype-specific gating. J Biol Chem 286, 8577–8584.

Garibaldi M, Fattori F, Riva B, Labasse C, Brochier G, Ottaviani P, Sacconi S, Vizzaccaro E, Laschena F, Romero NB, Genazzani A, Bertini E & Antonini G. (2016). A novel gain-of-function mutation in ORAI1 causes late-onset tubular aggregate myopathy and congenital miosis. Clin Genet 91, 780–786.

Hoth M & Penner R. (1993). Calcium release-activated calcium current in rat mast cells. J Physiol 465, 359–386.

Hou X, Pedi L, Diver MM & Long SB. (2012). Crystal Structure of the Calcium Release-Activated Calcium Channel Orai. Science 338, 1308–1313.

Lacruz RS & Feske S. (2015). Diseases caused by mutations in ORAI1 and STIM1. Ann N Y Acad Sci 1356, 45–79.

Lee KP, Yuan JP, Zeng W, So I, Worley PF & Muallem S. (2009). Molecular determinants of fast Ca^2+^-dependent inactivation and gating of the Orai channels. Proc Natl Acad Sci U S A 106, 14687–14692.

Lis A, Peinelt C, Beck A, Parvez S, Monteilh-Zoller M, Fleig A & Penner R. (2007). CRACM1, CRACM2, and CRACM3 are store-operated Ca^2+^ channels with distinct functional properties. Curr Biol 17, 794–800.

McNally BA, Somasundaram A, Yamashita M & Prakriya M. (2012). Gated regulation of CRAC channel ion selectivity by STIM1. Nature 482, 241–245.

Mullins FM & Lewis RS. (2016). The inactivation domain of STIM1 is functionally coupled with the Orai1 pore to enable Ca2+-dependent inactivation. J Gen Physiol 147, 153–164.

Mullins FM, Park CY, Dolmetsch RE & Lewis RS. (2009). STIM1 and calmodulin interact with Orai1 to induce Ca^2+^-dependent inactivation of CRAC channels. Proc Natl Acad Sci USA 106, 15495–15500.

Mullins FM, Yen M & Lewis RS. (2016). Orai1 pore residues control CRAC channel inactivation independently of calmodulin. J Gen Physiol 147, 137–152.

Navarro-Borelly L, Somasundaram A, Yamashita M, Ren D, Miller RJ & Prakriya M. (2008). STIM1-Orai1 interactions and Orai1 conformational changes revealed by live-cell FRET microscopy. J Physiol 586, 5383–5401.

Neher E. (1986). Concentration profiles of intracellular calcium in the presence of a diffusible chelator. . Experimental Brain Research Series 14, 80–96.

Nesin V, Wiley G, Kousi M, Ong EC, Lehmann T, Nicholl DJ, Suri M, Shahrizaila N, Katsanis N, Gaffney PM, Wierenga KJ & Tsiokas L. (2014). Activating mutations in STIM1 and ORAI1 cause overlapping syndromes of tubular myopathy and congenital miosis. Proc Natl Acad Sci U S A 111, 4197–4202.

Palty R, Stanley C & Isacoff EY. (2015). Critical role for Orai1 C-terminal domain and TM4 in CRAC channel gating. Cell Res 25, 963–980.

Park CY, Hoover PJ, Mullins FM, Bachhawat P, Covington ED, Raunser S, Walz T, Garcia KC, Dolmetsch RE & Lewis RS. (2009). STIM1 clusters and activates CRAC channels via direct binding of a cytosolic domain to Orai1. Cell 136, 876–890.

Prakriya M & Lewis RS. (2006). Regulation of CRAC channel activity by recruitment of silent channels to a high open-probability gating mode. J Gen Physiol 128, 373–386.

Prakriya M & Lewis RS. (2015). Store-Operated Calcium Channels. Physiol Rev 95, 1383–1436.

Scrimgeour N, Litjens T, Ma L, Barritt GJ & Rychkov GY. (2009). Properties of Orai1 mediated store-operated current depend on the expression levels of STIM1 and Orai1 proteins. J Physiol 587, 2903–2918.

Srikanth S, Jung HJ, Ribalet B & Gwack Y. (2010). The intracellular loop of Orai1 plays a central role in fast inactivation of Ca2+ release-activated Ca2+ channels. J Biol Chem 285, 5066–5075.

Stern MD. (1992). Buffering of calcium in the vicinity of a channel pore. Cell Calcium 13, 183–192.

Word JM, Lovell SC, LaBean TH, Taylor HC, Zalis ME, Presley BK, Richardson JS & Richardson DC. (1999). Visualizing and quantifying molecular goodness-of-fit: small-probe contact dots with explicit hydrogen atoms. J Mol Biol 285, 1711–1733.

Yamashita M, Navarro-Borelly L, McNally BA & Prakriya M. (2007). Orai1 mutations alter ion permeation and Ca2+-dependent fast inactivation of CRAC channels: evidence for coupling of permeation and gating. J Gen Physiol 130, 525–540.

Yamashita M, Somasundaram A & Prakriya M. (2011). Competitive modulation of CRAC channel gating by STIM1 and 2-aminoethyldiphenyl borate (2-APB). J Biol Chem 286, 9429–9442.

Yamashita M, Yeung PS, Ing CE, McNally BA, Pomes R & Prakriya M. (2017). STIM1 activates CRAC channels through rotation of the pore helix to open a hydrophobic gate. Nat Commun 8, 14512.

Yeung PS, Yamashita M, Ing CE, Pomes R, Freymann DM & Prakriya M. (2018). Mapping the functional anatomy of Orai1 transmembrane domains for CRAC channel gating. Proc Natl Acad Sci U S A 115, E5193–E5202.

Yeung PS, Yamashita M & Prakriya M. (2020). Molecular basis of allosteric Orai1 channel activation by STIM1. J Physiol 598, 1707–1723.

Zal T & Gascoigne NR. (2004). Photobleaching-corrected FRET efficiency imaging of live cells. Biophys J 86, 3923–3939.

Zhang SL, Yeromin AV, Hu J, Amcheslavsky A, Zheng H & Cahalan MD. (2011). Mutations in Orai1 transmembrane segment 1 cause STIM1-independent activation of Orai1 channels at glycine 98 and channel closure at arginine 91. Proc Natl Acad Sci USA 108, 17838–17843.

Zhang X, Pathak T, Yoast R, Emrich S, Xin P, Nwokonko RM, Johnson M, Wu S, Delierneux C, Gueguinou M, Hempel N, Putney JW, Jr., Gill DL & Trebak M. (2019). A calcium/cAMP signaling loop at the ORAI1 mouth drives channel inactivation to shape NFAT induction. Nat Commun 10, 1971.

Zhou Y, Cai X, Loktionova NA, Wang X, Nwokonko RM, Wang X, Wang Y, Rothberg BS, Trebak M & Gill DL. (2016). The STIM1-binding site nexus remotely controls Orai1 channel gating. Nat Commun 7, 13725.

Zweifach A & Lewis RS. (1993). Mitogen-regulated Ca^2+^ current of T lymphocytes is activated by depletion of intracellular Ca^2+^ stores. Proc Natl Acad Sci U S A 90, 6295–6299.

Zweifach A & Lewis RS. (1995a). Rapid inactivation of depletion-activated calcium current (I_CRAC_) due to local calcium feedback. J Gen Physiol 105, 209–226.

Zweifach A & Lewis RS. (1995b). Rapid inactivation of depletion-activated calcium current (ICRAC) due to local calcium feedback. The Journal of general physiology 105, 209–226.

